# Spatial navigation through evolution: a single-cell atlas of the mammalian entorhinal cortex

**DOI:** 10.64898/2026.05.07.723541

**Authors:** Dorottya Maria Ralbovszki, Jesse J. Westfall, Yuki Mori, Stamatios A. Tahas, Mads F. Bertelsen, Irina Korshunova, Josefine Skov, Jan Gorodkin, Martin Hemberg, Stefan E. Seemann, Vanessa Hall, Menno P. Witter, Konstantin Khodosevich

**Affiliations:** Biotech Research and Innovation Centre (BRIC), Faculty of Health and Medical Sciences, University of Copenhagen, 2200 Copenhagen, Denmark; Section for Parasitology and Pathobiology, Department of Veterinary and Animal Sciences, Faculty of Health and Medical Sciences, University of Copenhagen, 1870 Frederiksberg, Denmark; Core Facility for Integrated BioImaging, Center for Core Facilities, University of Copenhagen, Copenhagen, Denmark; Copenhagen Zoo, Copenhagen, Denmark; Preclinical Disease Biology, Department of Veterinary and Animal Sciences, Faculty of Health and Medical Sciences, University of Copenhagen, 2200 Copenhagen, Denmark; Gene Lay Institute of Immunology and Inflammation, Brigham and Women’s Hospital, Massachusetts General Hospital and Harvard Medical School, Boston, USA; Magle Chemoswed AB, Agneslundsvägen 27, 212 15 Malmö, Sweden; Kavli Institute for Systems Neuroscience, Faculty of Medicine and Health Sciences, Norwegian University of Science and Technology, Trondheim, Norway

## Abstract

Spatial navigation is a fundamental mammalian ability, supported by the entorhinal cortex (EC), a structurally conserved yet functionally diverse region across mammalian species. However, how molecular signaling underlies both shared and species-specific navigational strategies remains unclear. Here, we present a cross-species single-cell atlas of the EC from human, Hamadryas baboon, mouse, and Egyptian fruit bat - species spanning distinct evolutionary lineages and navigational demands, including true 3D navigation in bats. Using this resource, we identify conserved principal neuron populations as well as species-specific innovations, including mixed-layer or functional identities and fruit bat-specific subtypes. GABAergic interneurons neurons show strong conservation of somatostatin (SST) and parvalbumin (PV) families, while VIP GABAergic neurons exhibit pronounced species-specific divergence, with an expanded repertoire in primates. Integration with whole-brain diffusion tensor imaging reveals conserved and species-specific connectivity between the EC, hippocampus, and sensory cortices. Major species-specific cellular innovations were further validated using orthogonal histological approaches, confirming their anatomical and laminar organization. Overall, this atlas provides a comparative framework available for the research community to dissect the molecular, cellular, and circuit principles underlying conserved and specialized spatial navigation across mammals.

## Introduction

Spatial navigation encompasses the cognitive processes that allow animals to determine and reach one location from another, ranging from local exploration to long-distance migration across thousands of kilometers. It is a fundamental function of the brain in mobile animals, requiring perception of the environment, and the formation and recall of internal representations. In general, mammals rely on diverse sensory modalities to navigate, including vision, olfaction, echolocation, and magnetoreception [1–4]. Most mammals, including primates, navigate primarily using remembered landmarks [3,4].

Spatial navigation has been studied across a wide range of mammalian species. Although most neuroanatomical, electrophysiological, and behavioral studies have focused on rats and monkeys [5], bats have also been extensively studied as the only mammals capable of powered flight [6], introducing navigation in three dimensions. Bats exhibit remarkable navigation abilities, relying on a combination of sensory modalities. Echolocating bats emit sound pulses and analyze returning echoes to estimate distance, direction, velocity, size, and texture of objects in their surroundings. Additionally, Old World fruit bats possess exceptional low-light vision, with visual acuity that surpasses that of humans under night-like conditions [2].

It has been established that the entorhinal cortex (EC), together with the hippocampus, plays a crucial role in spatial navigation [1]. The EC provides the main cortical input to the hippocampus and forms a processing system connecting the hippocampus and several cortical areas [7]. The EC is primarily divided into the medial and lateral EC (MEC and LEC, respectively), which are found to differ in their functional roles, connectivity, and anatomical features in all studied species [5,7,8]. The MEC is primarily implicated in processing spatial information such as direction, speed, and grid-based location signals, while the LEC plays a role in object recognition.

Four principal neuronal types defined by their electrophysiological properties underpin spatial representation within the hippocampal formation: grid cells, place cells, border cells, and head direction cells [1]. While place cells are restricted to the hippocampus, the other cell types are found in the EC and other brain regions [9]. These neuronal types that coordinate spatial navigation were first identified in rats and mice. Subsequently, all four cell types were described in bats [2,10]. In primates, grid, border and head direction cells were identified in the EC [9], while in the hippocampus, primate-specific view cells were reported [11].

Single-cell genomics has become a key tool for elucidating cell taxonomy and the evolutionary architecture of the brain. Cross-species integration has revealed conserved brain cell types while capturing transcriptomic divergence across rodents, primates and other species [12–15]. Here, we present a single-cell atlas of the EC from four mammalian species: human, Hamadryas baboon, mouse, and Egyptian fruit bat. Constructing a single-cell atlas across four evolutionarily distant mammals including model and non-model species, enables the identification of evolutionarily conserved cell populations within the spatial processing system, assessment of cell type-specific transcriptomic divergence, and linking molecular identity to known anatomical and functional roles. To further bridge gene expression with brain architecture, we investigated MEC connectivity in baboons and fruit bats using diffusion tensor magnetic resonance imaging (DTI). Together, this atlas provides a unique resource, which can be implemented by the field for investigating the transcriptomic foundations of spatial navigation and memory across mammals and instruct further mechanistic functional studies.

## Results

### Constructing reference genomes of non-model species and mapping orthologs across mammals

To produce molecular-to-connectivity atlas of spatial navigation in mammals, we first performed single-nucleus RNA-sequencing (snRNA-seq) analysis capturing the transcriptional conservation and divergence in the EC across mammals. We chose two primate species – human (*Homo sapiens*) and Hamadryas baboon (*Papio hamadryas*), one rodent – mouse (*Mus musculus*), and one bat – Egyptian fruit bat (*Rousettus aegyptiacus*) - species. The selection of species was guided by previous spatial navigation studies, different navigational strategies and our aim to investigate primate-specific gene orthologs or cell subtypes that might underlie spatial navigation. Mouse is the most used model organism with the best described spatial navigation, from molecular mechanisms to behavior. Including data from humans is essential because it reveals how conserved EC circuits are adapted by molecular and cellular specializations supporting complex human cognitive navigation, while also enabling the identification of vulnerable EC cell types that are selectively affected in human brain disorders, thereby improving both our understanding of human cognition and the translational relevance of disease models. To complement available human EC datasets, we collected a novel primate EC single-cell dataset from baboons. Hamadryas baboons live a terrestrial lifestyle within complex social hierarchies of up to 700 individuals and exhibit higher learning abilities compared to macaques, which live in smaller, arboreal groups [16]. Fruit bats are widely used to study 3D spatial navigation and use a unique navigational strategy: echolocation together with vision. To this end, we collected fresh postmortem entorhinal cortex samples from juvenile and subadult Hamadryas baboons and Egyptian fruit bats (Fig. 1A,B) and utilized previously published human and mouse snRNA-seq data [14,17].

**Figure 1.**
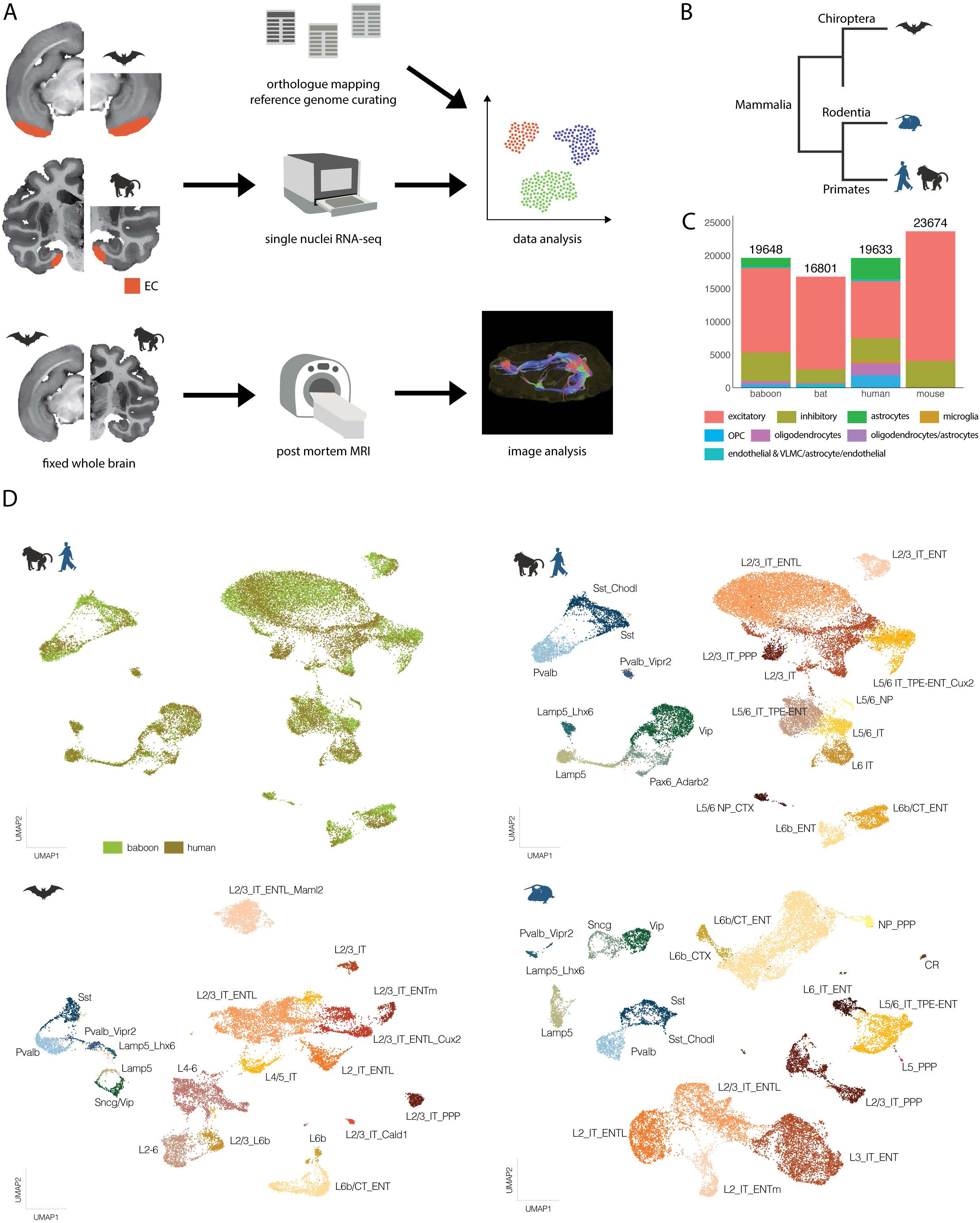
Cell type and neuronal subtype diversity in the EC at a single-cell resolution. **(A)** Overview of the experimental workflow, including sample collection and analysis steps. **(B)** Phylogenetic relationship of the four species analyzed. Previously published mouse and human datasets are indicated in blue. **(C)** Distribution of major cell types detected in each species. Numbers indicate total nucleus counts in the corresponding species. **(D)** UMAP visualizations of nuclei from human and baboon samples in the primate EC neuronal dataset (left) and neuronal subtypes annotated, colored and labeled by neuronal subtype (right). **(E)** UMAP visualizations of neuronal subtypes detected in the EC of fruit bats and mice, colored and labeled by neuronal subtype.

Importantly, the reference genomes of non-model target species in our atlas, namely, baboon and fruit bat, were not formatted for direct use in single-nucleus expression quantification. Since generating reference genomes compatible with single-cell transcriptomics workflow for non-model species is a crucial foundation for comparative single-cell studies, we constructed compatible reference genomes ready to be used by the Cell Ranger pipeline. In short, missing mitochondrial genes were added to the fruit bat reference genome, and reference genomes were filtered for protein-coding genes. The constructed reference genomes have been made publicly available as FASTA and GTF files (see Methods).

We performed snRNA-seq for the EC dissected from baboon (n=4) and fruit bat (n=4) using 10x Genomics 3’ kit v3.1. The reference genomes were applied to quantify gene expression at a single-nucleus resolution (Fig. 1A). We obtained average mapping rates of 64.7% for fruit bats and 82.2% for baboons, excluding one baboon sample with a 70.5% mapping rate that later failed preprocessing.

To compare gene expression across studied species, we constructed a gene ortholog database tailored to our expression data for the target species (Fig. 1A,B). In brief, pairwise orthologs between baboon-human, human-mouse, and fruit bat-mouse were extracted and filtered to retain genes that are expressed in our snRNA-seq data. In order to maximize the number of gene orthologs between our target species, genes with multiple orthologs were converted into one-to-one orthologs. Mapped genes across species represented approximately 90% of the aligned baboon genes (n = 18,535) and fruit bat genes (n = 16,744).

### Neuronal diversity in the entorhinal cortex in mammals

Following expression quantification and filtering, we retained 19,648 high-quality nuclei from baboons, 16,801 from fruit bats, 23,674 from mice, and 19,633 from humans (Fig. 1C). Through unsupervised clustering and analysis of marker gene expression patterns, we defined seven, five, and two major cell types in primates (humans and baboons), fruit bats, and mice (mouse dataset contained only neurons [17]), respectively. In total, we identified 55,096 excitatory neurons, 13,800 inhibitory neurons, 4,587 astrocytes, 2,250 oligodendrocytes, 2,469 OPCs, 399 microglia, and 438 endothelial cells (Figs. 1C, S1). As our primary focus in this study was to capture neuronal diversity contributing to spatial navigation processing in the EC across our target species, non-neuronal cells were filtered out. Human and baboon neurons were integrated into a primate EC dataset, which was used for subsequent neuronal annotation, facilitated by label transfer from previously constructed EC reference snRNA-seq dataset [17] (Fig. 1D).

We identified 12 excitatory neuron subtypes in primates and mice, and 14 in fruit bats (Fig. 1D,E). These excitatory subtypes were annotated based on layer-specific marker genes, axonal projection properties (isocortex [CTX], intratelencephalic [IT], near projecting [NP], cortical layer 6b [L6b]), and anatomical area (entorhinal [ENT] including lateral [ENTl] and medial [ENTm], para-post-pre-subiculum [PPP], temporal association-perirhinal-ectorhinal areas [TPE]), coupled with cluster specific marker genes. Prediction of these properties was partially established by label transfer from reference EC snRNA-seq dataset [17] that contained information about projection, layer and anatomical location. Entorhinal superficial and deep layer subtypes appeared in all species, with L2_ENT subtypes being absent in primates. L2/3_IT_ENTL, L2/3_IT_PPP, and L6b/CT subtypes were detected in all species. For further subdivision of neuron subtypes with the same layer-projection-area, we included a cluster specific marker gene in the final label.

We identified two L5/6_IT_TPE-ENT annotated clusters in primates. Interestingly, one expressed the established superficial layer marker *Cux2* and neighbored superficial layer clusters (termed L5/6_IT_TPE-ENT_Cux2), while the other did not express *Cux2* and clustered together with other deep layer subtypes (termed L5/6_IT_TPE-ENT) (Figs. 1D, S2A).

Fruit bats presented multiple clusters with the same superficial layer-projection-area such as L2/3_IT (cortical layer II and III neurons that project to intratelencephalic areas) and L6b/CT_ENT (cortical layer VIb/corticothalamic neurons that project intratelencephalically). Furthermore, these clusters could all be distinguished by marker genes that were specific to these clusters - *Maml2* and *Cux2* for L2/3_IT_ENTL, and *Cald1* for L2/3_IT (Figs. 1E, S2B,D). Finally, we found two excitatory neuron clusters where the gene expression pattern could not be annotated to a specific layer location – L2-6 and L2/3_L6b (Fig. 1E), emphasizing limited knowledge in layer-specific marker expression for fruit bats.

We detected 8 inhibitory neuron subtypes in primates and in mice, whereas in fruit bats we could identify only 6 subtypes (Fig. 1E). In each species, inhibitory neuron subtypes displayed robust expression of canonical marker genes. Thus, the differential developmental origin was identified by lineage markers, such as *SoxC* and *LhxC* for medial ganglionic eminence (MGE)- derived neurons across all species, and *Adarb2* and *Prox1* for caudal ganglionic eminence (CGE)-derived neurons in primates and mice. Interestingly, *Prox1* expression was not detectable in *Gad1+, Sox6-* neurons of fruit bats, while *Adarb2* marker could not be used, since fruit bats lack its orthologous gene. Thus, we used *Vip*, *Reln* and *Cnr1* markers to determine CGE-derived neurons in fruit bats (Fig. S2D).

Two subtypes of Pvalb (Pvalb and Pvalb_Vipr2) and Lamp5 (Lamp5 and Lamp5_Lhx6) neurons were identified in all species (Fig. 1D,E). The Sst subtype was clearly detected by strong *Sst* expression in all species, whereas *Npy+* Sst_Chodl long-range projection inhibitory neurons were found in primates and mice, but absent in bats. As expected, mice had additionally Vip and Sncg neuronal subtypes (Fig. 1E). Based on *Vip* expression and cell type labels provided by the reference EC dataset, presumable counterparts of Vip and Sncg inhibitory neuron subtypes formed one joint cluster in fruit bats (thus named Sncg/Vip subtype). In primates, we identified Vip subtype and the primate homolog of Sncg subtype - Pax6_Adarb2, using previously established markers [12].

### Cross-species integration reveals conserved neuronal subtypes in the mammalian entorhinal cortex

We next investigated conserved cell types in the EC across mammals by integrating the transcriptomic datasets from primates, fruit bats, and mice, thus producing a single-cell atlas of the EC. In total, we detected 28 neuronal subtypes: 10 were species-enriched, and 3 lacked marker expression for assigning them specifically to superficial or deep layers (Fig. 2A). 23 subtypes included nuclei from all four species (Fig. 2B). A subtype was classified as species-enriched if a single species contributed more than 75% of the nuclei within that cluster; for this analysis, primates were considered jointly. The 10 species-enriched subtypes included two fruit bat-enriched excitatory superficial-layer neuron subtypes (L2/3_IT_ENTL_bat, L2/3_IT_ENTL_Maml2_bat), three mouse- and one primate-enriched excitatory deep-layer neuron subtypes (L5_PPP_mouse, L6b/CT_ENT_mouse, L6_IT_ENTL_mouse, L6_IT_primate), two mixed-layer neuron subtypes enriched by primates and fruit bats (L2-6_bat, L2-6_primate), and one primate- and one mouse-enriched inhibitory subtype (Vip_primate, Sst_Chodl_mouse) (Fig. 2A).

**Figure 2.**
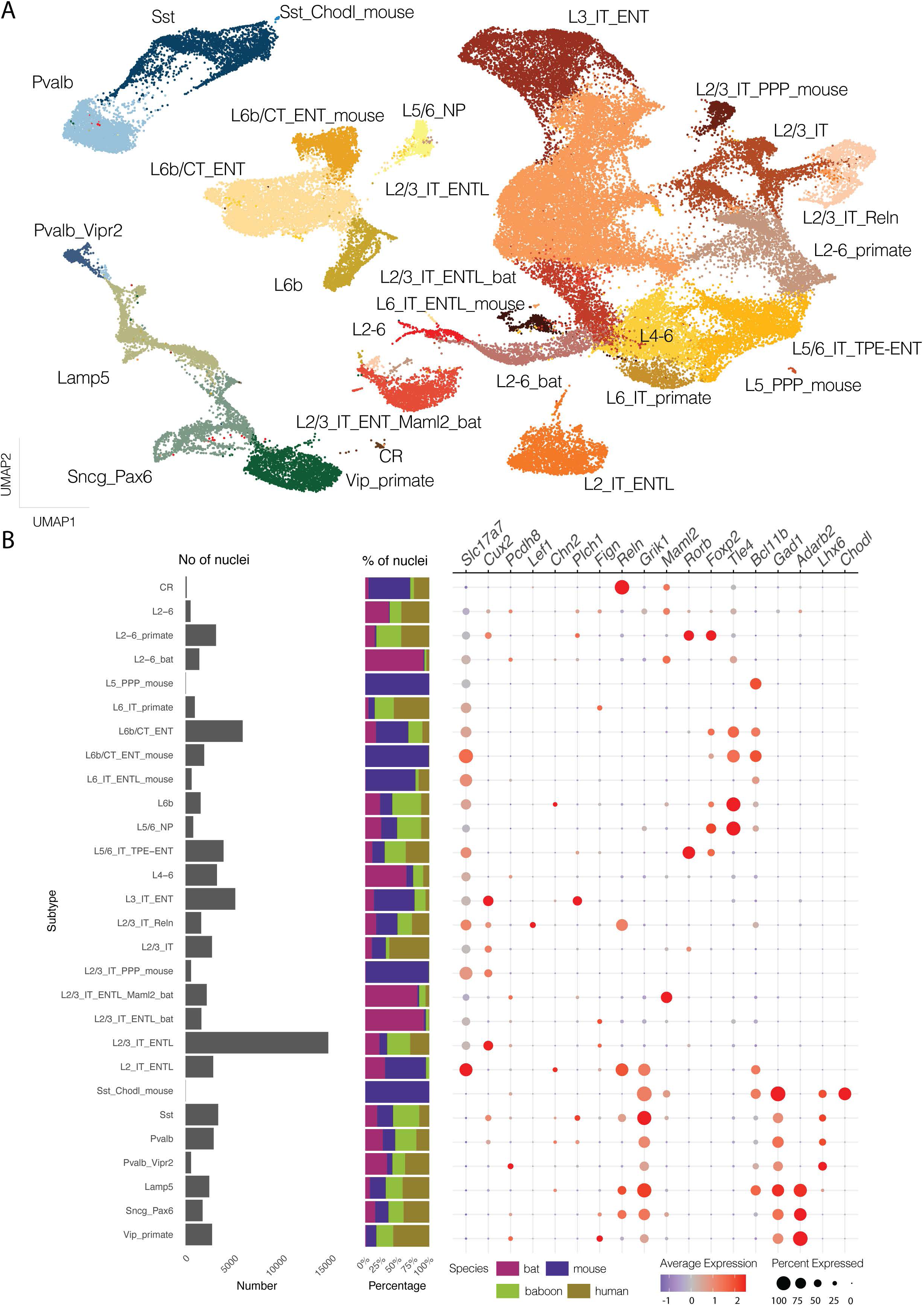
A cross-species atlas of neuronal subtypes in the entorhinal cortex. **(A)** UMAP visualization of neurons in the entorhinal cortex following cross-species integration, annotated by neuronal subtype. **(B)** Bar plots showing the number of nuclei (left) and relative species contributions (middle) for each subtype. Dot plot (right) displays expression of selected marker genes across identified subtypes.

To better understand the evolutionary dynamics of gene expression conservation, we assessed transcriptional divergence across species for each shared subtype. Divergence patterns reflected evolutionary distances: it was lowest between primates and highest between mice and fruit bats (Figs. 3A, S3A, see evolutionary relations in Fig. 1B). Subtypes with relatively few nuclei or strong species bias (*i.e.*, predominantly found in one species) showed higher transcriptomic divergence, likely due to increased variance from limited sampling, such as Cajal-Retzius cells (CR) and L2/3_IT_ENTL_bat (Fig. 3A).

**Figure 3.**
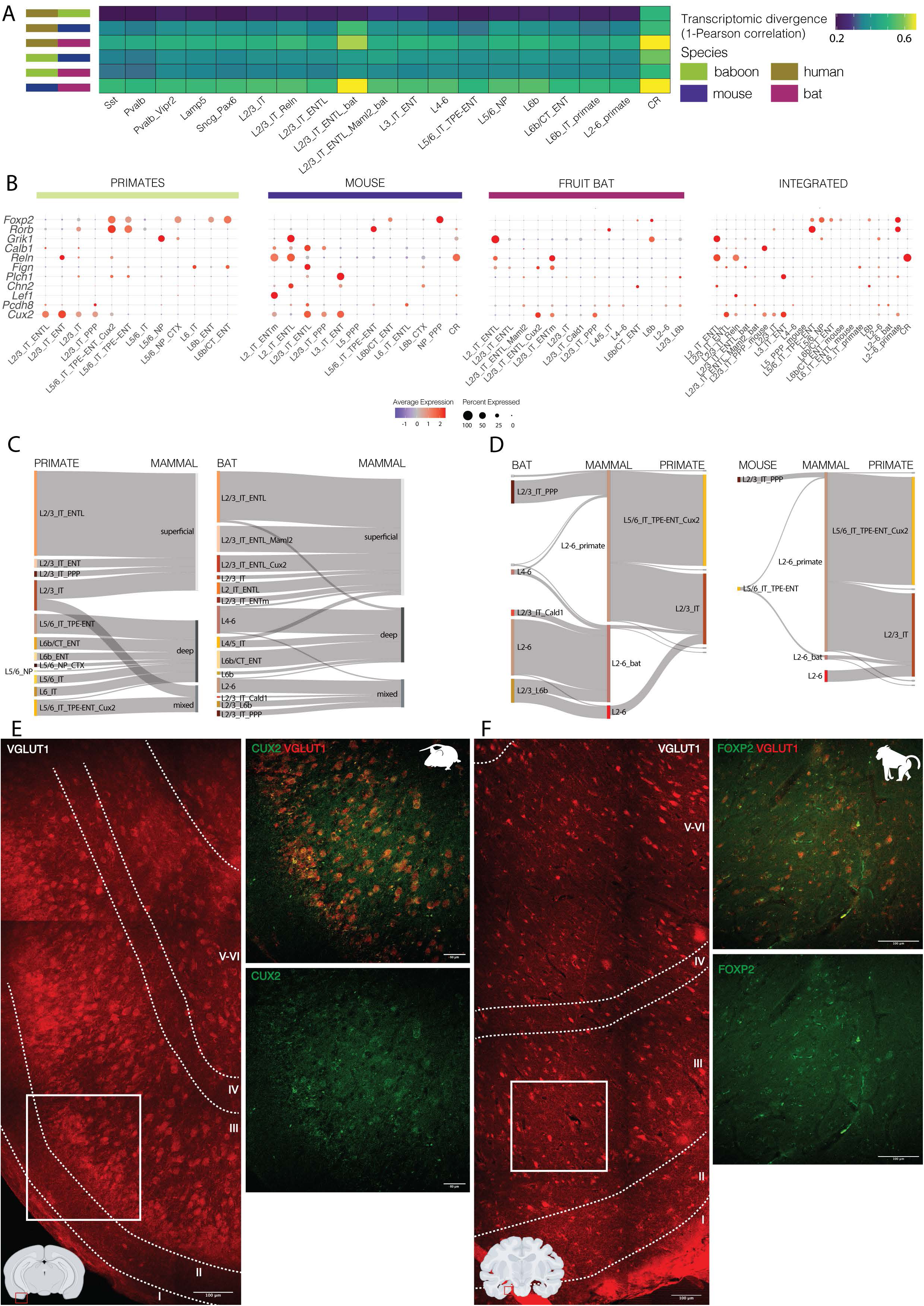
Divergent features of excitatory neurons in the entorhinal cortex across mammalian species. **(A)** Raw transcriptomic divergence scores across shared neuronal subtypes reflect evolutionary distances. **(B)** Sankey plot shows transitions between species-specific annotations (left side) and cross-species subtypes grouped by predicted cortical layer identity (right side). **(C)** Dot plot showing expression of EC-specific marker genes across excitatory subtypes in species-specific datasets and the integrated dataset (right). These markers were reported in mice. Additionally, the expression of mid and deep layer markers *Rorb* and *Foxp2* respectively are also shown. **(D)** Detailed Sankey plot highlighting the origin of neurons contributing to mixed-layer subtypes. **(E)** Fluorescent immunohistochemistry staining of the mouse and baboon MEC. Mouse MEC was stained against glutuminergic marker VGLUT1 and superficial layer marker CUX2. Baboon MEC was stained against glutuminergic marker VGLUT1 and excitatory layer marker FOXP2.

To further investigate species-specific divergence within shared subtypes, differentially expressed genes (DEGs) were identified between species within each neuronal subtype. Surprisingly, we found strong pan-neuronal species-specific transcriptional signatures that dominated over neuronal subtype-specific gene expression. We illustrate this pan-neuronal transcriptional signature by showing species-specific DEGs captured in two distant neuronal subtypes - the largest excitatory neuron subtype, L2/3_IT_ENTL, and a well-conserved inhibitory neuron subtype, Pvalb (Fig. S3B). Thus, while some established subtype-specific marker genes are conserved across mammals (*e.g.*, *Cux2* or *Pvalb*), there is a strong domination of pan-neuronal species-specific transcriptional signature reflected in DEGs across all neuronal cell types. Highly variable genes between species (humans, baboons, fruit bats, and mice) effectively captured this species-level divergence shared by all neuronal subtypes (Fig. S3B).

### Conservation and species specialization across layers of excitatory neurons of the entorhinal cortex

Excitatory neurons largely maintained clustering by predicted layer origin in the cross-species integration. Subtypes annotated in species-specific datasets mostly retained their layer identity (Fig. 3C). Neurons annotated as L2/3_IT_ENTL and L3_IT_ENT accounted for over 35% of all excitatory neurons in the cross-species integrated dataset. Conserved superficial-layer subtypes were characterized by canonical superficial layer markers *Cux2* and/or *Pcdh8* (Figs. 2B, 3B). Entorhinal markers reported in mice [17] were also detected across various ENT and superficial-layer annotated subtypes in the cross-species integrated dataset. The non-primate *Cux2*-negative L2_IT_ENTL subtype was marked by *Chn2*, *Grik1*, and *Reln* (Figs. 2B, 3B), likely corresponding to pyramidal, fan, and multimodal cells [7]. The layer II intratelencephalic MEC marker *Lef1* was expressed in the L2/3_IT_Reln subtype marking stellate and pyramidal cells (Figs. 2B, 3B), and the L3_IT_ENT subtype expressed layer III MEC marker *Plch1* where pyramidal and multipolar cells are found [7]. All subtypes with predicted ENTL location expressed superficial layer LEC marker *Fign*, except for the L2/3_IT_ENTl_Maml2_bat subtype (Figs. 2B, 3B). When we looked at these entorhinal markers in species-specific datasets, fruit bats expressed most of these ENT markers, although they did not exactly follow the expression pattern found in mice (Fig. 3B). In primates, these markers were not prominent in superficial EC subtypes; *Plch1* was expressed in the L2/3_IT subtype, and the L2/3_IT_ENT subtype was *Reln+* (Fig. 3B).

Deep-layer subtypes comprised 27% of total neurons and formed several conserved and species-specific clusters. Conserved clusters included three L6b annotated neuronal subtypes, which consistently expressed canonical deep-layer markers *Tle4* and *Bcl11b* (Figs. 2B, 3A, S4A). Another conserved subtype was annotated as L5/6_NP excitatory neurons, which expressed *Grik1* and *Kcnip1*, known markers of NP neurons [17] (Figs. 2B, S2E). Among species-enriched deep-layer subtypes, we detected two mouse-enriched: L6b/CT_ENT_mouse and L5_PPP_mouse (Fig. 2B). Some EC-specific superficial markers were expressed in deep layer subtypes in the primate-, and fruit bat-specific datasets (Fig. 3B). In primates, *Plch1* was detected in both L5/6_IT_TPE-ENT and L5/6_IT_TPE-ENT_Cux2 subtypes, and *Fign* was expressed in the L6b/CT_ENT subtype (Fig. 3B). The layer II intratelencephalic LEC marker *Chn2* was expressed in the L6b/CT_ENT subtype in fruit bats (Fig. 3B).

Interestingly, while most excitatory subtypes showed clear layer-specific expression patterns following cross-species integration, three subtypes (L2–6, L2–6_bat, and L2–6_primate) could not be annotated to either deep or superficial layers and were therefore referred to as mixed-layer subtypes (Figs. 2; S4B). Since we used mouse EC dataset to aid annotation, the low number of mouse neurons contributing to mixed layer clusters is expected (Figs. 3C; S4C). Additionally, fruit bat excitatory neurons with unresolved laminar identity may express a combination of layer-markers unique to fruit bats. Alternatively, in species-enriched mixed subtypes, species-specificity may override canonical layer marker expression. The two mixed layer excitatory subtypes in the fruit bat dataset consistently mapped to the three mixed-layer subtypes in the cross-species integrated dataset (Fig. 3C,D). The fruit bat-enriched L2–6_bat subtype co-expressed deep-layer marker *Tle4*, the superficial marker *Pcdh8*, and fruit bat-specific marker genes *Maml2* and *Zbtb20* (Figs. 2B, S2F). Neurons contributing to this subtype primarily originated from the two mixed-layer subtypes identified in fruit bats (L2–6 and L2/3_L6b) with minimal input from primates (Fig. 3C,D). These fruit bat subtypes also showed expression patterns similar to L2-6_bat subtype (Figs. 3B, S2D) highlighting a marker gene combination unique to fruit bats.

Notably, some excitatory neurons that were initially annotated as either superficial- or deep-layer–specific in primates shifted into mixed-layer subtypes following cross-species integration (Fig. 3C,D). Further analysis of these mixed-layer neurons revealed that the L2– 6_primate subtype co-expressed superficial (*Cux2*), mid (*Rorb*), and deep-layer (*Foxp2*) markers (Figs. 2B, 3B, S4B). Interestingly, although the L2–6_primate subtype also contained neurons from mouse and fruit bat, these cells did not express either mid- (*Rorb*) or deep-layer (*Foxp2*) markers (Figs. 3B-D, S2B,C). Instead, mouse and fruit bat neurons within this cluster were predominantly annotated as superficial-layer neurons and were marked by *Cux2* expression (Figs. 3B-D; S2B,C), which we validated in mouse MEC by immunohistochemistry (Fig. 3E). Tracing L2-6_primate neurons back to the primate-only dataset revealed their origin from *Cux2*-expressing neuronal subtypes in both superficial- (L2/3_IT) and deep-layers (L5/6_IT_TPE-ENT_Cux2) (Fig. 3C,D). Although *Cux2* is classically considered a marker of superficial cortical layers in mammals, its expression in deeper layers has previously been reported in primate granular dorsolateral prefrontal cortex [12]. Likewise, *Rorb* and *Foxp2* are generally considered mid- and deep-layer markers, respectively, in mammals [17–19], whereas primate-specific mid-layer expression of *Foxp2* was reported in the dorsolateral prefrontal cortex [12]. Our data confirms separation of *Cux2* superficial- vs *Rorb/Foxp2* deep-layer expression in mouse and bat EC (Figs. 3B, S2B,C). In contrast, primate EC exhibited additional superficial-to-mid-layer expression of *Rorb* and *Foxp2* (Figs. 3B, S2A). To validate such extended expression of deep markers into superficial layers, we labeled FOXP2+ neurons by immunohistochemistry in baboon MEC, which showed that FOXP2+ neurons are present from layer VI to layer III (Fig. 3E). Together, these findings highlight the complexity of assigning laminar identity in cross-species single-cell datasets and underscore substantial species-specific expression differences even of classical layer-specific markers, which suggest differences in layer-specialization across mammalian EC.

### Inhibitory neuron subtypes are conserved across mammals in the entorhinal cortex

Opposite to excitatory neurons, inhibitory neurons did not form new clusters in cross-species integrated datasets that were not present in the species-specific datasets (Fig. 4A). Canonical markers clearly delineated CGE- and MGE-derived inhibitory subtypes across species. In MGE, *SoxC* and *LhxC* were expressed in all species (Fig. 4B). Likewise, *Prox1* was pan-CGE across all species, whereas the other mammalian CGE-lineage markers, *Adarb2* and *Nr2f2*, were absent in fruit bats (Figs. 2B, 4B).

**Figure 4.**
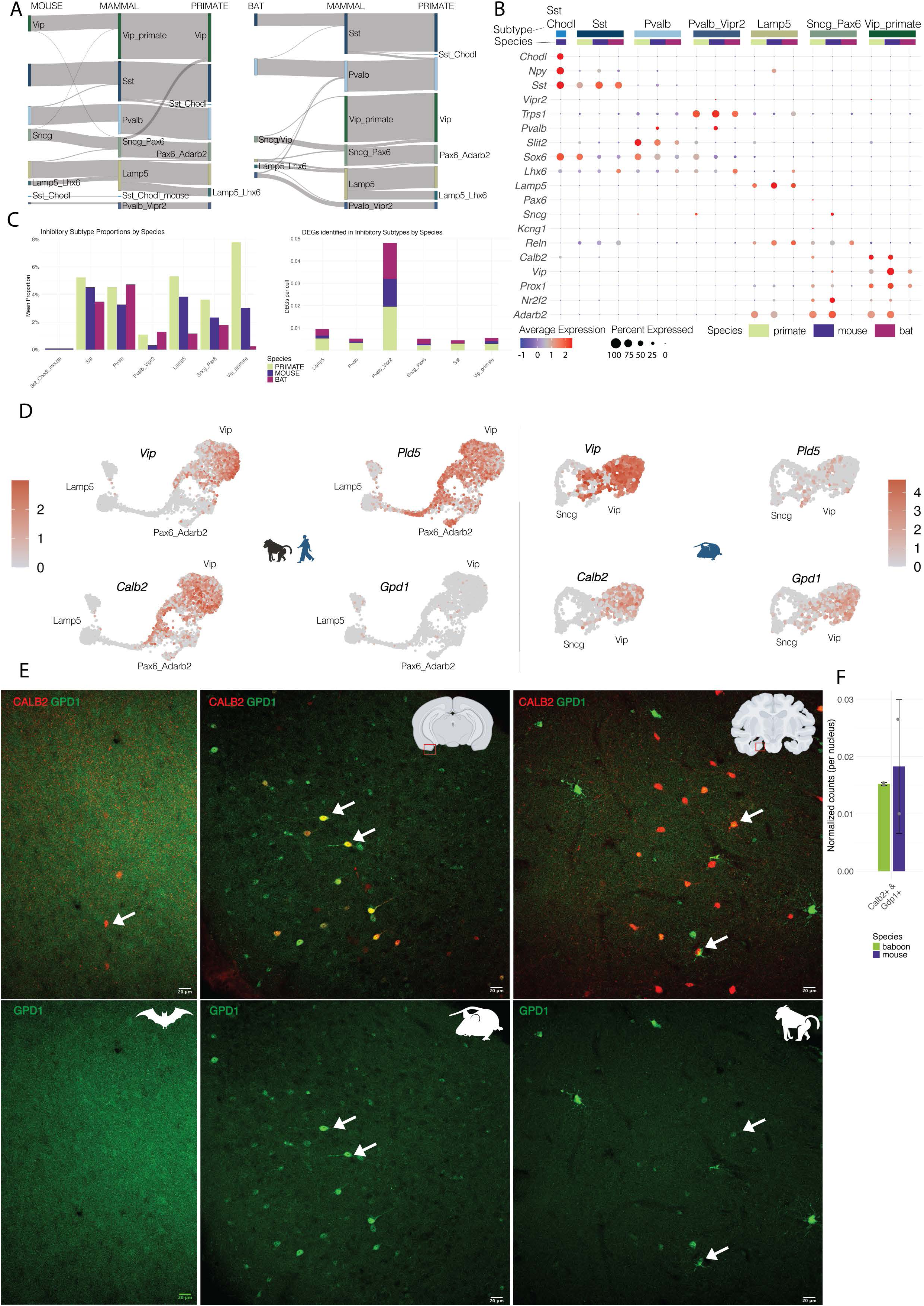
Inhibitory neuronal subtypes are conserved across species with species-specific gene expression signatures. **(A)** Sankey plot showing the correspondence between inhibitory neuronal subtypes annotated in individual species and those identified in the cross-species integrated dataset. **(B)** Dot plot showing the expression of canonical inhibitory marker genes across cross-species subtypes and species. **(C)** Relative proportions of inhibitory subtypes among all neuronal subtypes are shown for each species, with humans and baboons grouped together as primates. To the right, the bar plot shows the number of DEGs per nuclei identified within species in each inhibitory subtype. Subtypes such as Pvalb_Vipr2 and Lamp5 exhibit the highest number of DEGs. **(D)** UMAPs of primate and mouse *Vip+* and *Calb2+* inhibitory neurons highlighting species-specific expression of selected species-specific markers such as *Pld5* (primate), and *Gpd1* (mouse). **(E)** Representative images of validation of CALB2 and GPD1 positive neurons in fruit bat, mouse and baboon MEC. Arrows indicate example double positive neurons (except in the case of fruit bats where we only detected CALB2+ neurons). **(F)** Number of Calb2+, Gpd1+ positive neurons visualized on a bar plot. The neurons were identified during immunofluorescence staining of mouse and baboon MEC.

As mentioned before, transcriptomic profiles for Vip and Sncg neurons in fruit bats did not resolve into distinct subtypes and instead formed a shared group (Vip/Sncg) (Fig. 1D). However, cross-species integration allowed to split Vip/Sncg fruit bat neurons into Sncg (most neurons) and Vip subtypes (Fig. 4A). Notably, the Sncg_Pax6 subtype was the most heterogeneous and lacked a universal marker gene shared across all species (Fig. 4B). Nevertheless, the predicted homology between mouse Sncg and primate Pax6_Adarb2 subtypes was supported by their integration in the cross-species dataset (Fig. 4A).

While overall inhibitory subtypes were conserved, their relative proportions differed across species. As previously reported, CGE-derived inhibitory neurons are more abundant in primates [20], and we captured the same trend, which was particularly strong for Vip neurons (Figs. 4C, S4D). To identify genes that might underlie this expansion of Vip neurons in primates, we searched for species-specific gene expression and identified *Pld5* gene to be specific to primates, while *Gpd1* was enriched in mice (Figs. 4D, S4E). To validate prevalence of Vip/Gpd1 co-expression in mice and primates, we performed immunohistochemistry for a common marker for Vip neurons (CALB2) as well as for GPD1 in mouse and baboon MEC. We found CALB2/GDP1 double-positive neurons in both species, and these seemed more abundant in mice than in primates on representative sections (Fig. 4E,F).

Another interesting species-specific feature of Vip neurons was the complete lack of *Gpd1* gene expression in bats (Fig. S4E). Furthermore, bat’s CGE-derived neurons did not express the *Calb2* gene, which is considered to be general marker of Vip neurons in mammals [21] (Fig. 4B). To validate these findings at protein level, we labeled CALB2 and GPD1 in fruit bat MEC and although we found few CALB2+ neurons (which were likely principal neurons, based on fruit bat snRNA-seq data), we confirmed complete lack of GPD1 in bat’s MEC (Fig. 4E).

MGE-derived Pvalb and Sst subtypes were more conserved than CGE-derived and had comparable proportions across primates, mice and fruit bats (Figs. 4C, S2B). Interestingly, when subdividing primates, baboons exhibited surprisingly high proportions of both subtypes, in particular Sst (Fig. S2B). Pvalb and Sst subtypes also showed high concordance between their species-specific and cross-species integrated annotations (Fig. 4A). For instance, the Pvalb_Vipr2 subtype (putative chandelier cells) was robustly preserved across species and included almost exclusively previously Pvalb_Vipr2 labeled neurons (Fig. 4A). Despite forming a tight cluster in the cross-species integrated dataset (Fig. 2A), this subtype exhibited the highest number of DEGs (Fig. 4C). The Pvalb subtype (putative basket cells) showed consistent MGE-specific marker expression across species (Fig. 4B) and, together with the Sst subtype, exhibited low transcriptomic divergence (Fig. 3A), suggesting a high degree of conservation, consistent with one of its key roles, modulating grid cell activity in the MEC [22].

Sst_Chodl subtype represents unique long-range GABAergic neurons [23], which were recently shown to be particularly affected by neuropsychiatric risk factors [24]. Interestingly, in the integrated dataset, primate *Npy+ Chodl+* neurons (putative Sst_Chodl subtype in primates) clustered with the Sst subtype, not with mouse Sst_Chodl neurons that remained distinct in the cross-species integrated space (Fig. 4A), which might indicate strong divergence of primate and mouse Sst_Chodl neurons.

In summary, while canonical marker gene expression highlights strong conservation of inhibitory neuron types across species, there are species-specific changes in abundance and molecular profiles of inhibitory subtypes.

### Connectivity of the medial entorhinal cortex in mammals with different spatial navigation strategies

While single-cell transcriptomics provides detailed insights into the molecular identity of neuronal populations, it does not directly capture how these cells are embedded within functional brain circuits. We therefore complemented our single-cell analysis with diffusion tensor magnetic resonance imaging (DTI) tractography to examine the organization of the spatial navigation network in baboons and fruit bats. Importantly, transcriptomic differences across species were most pronounced in neuronal populations involved in sensory integration and long-range communication, particularly in superficial excitatory neurons. We therefore asked whether these molecular specializations are reflected at the level of mesoscale connectivity within the spatial navigation network. Furthermore, we investigated whether the use of echolocation in addition to vision in navigation and terrestrial versus flying lifestyle could influence the connectivity of the spatial processing center. Target anatomical structures were the MEC, hippocampus, visual cortex, and auditory cortex (Figs. 5A,B, S5-6).

**Figure 5.**
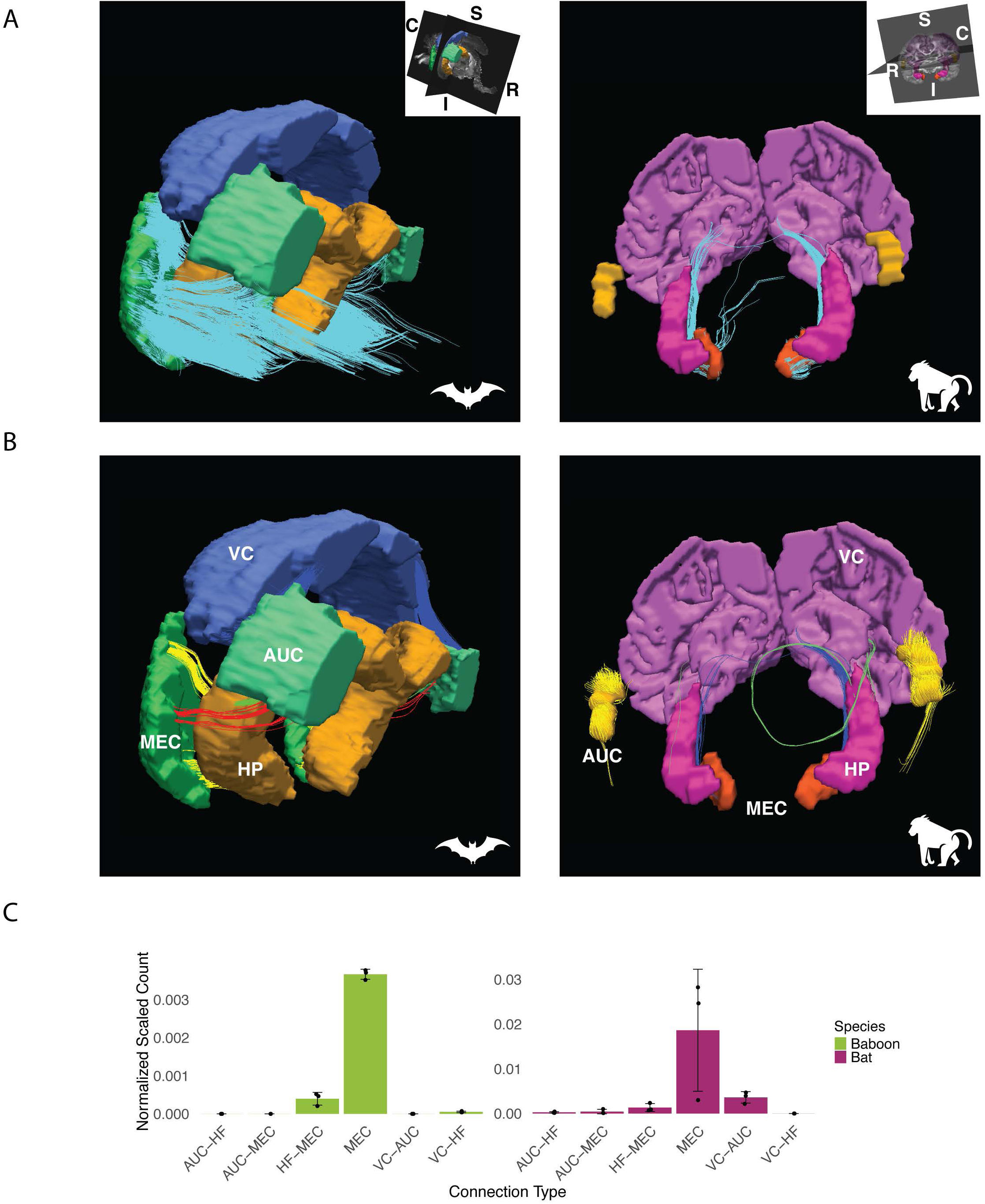
Connectivity of the MEC in the baboon and fruit bat. **(A)** Tractography-based connectivity (turquoise) of the MEC in fruit bats (left) and baboons (right). Anatomical planes: R - rostral, C - caudal, S - superior, I - inferior. **(B)** Tractography-based connectivity of the medial entorhinal cortex (MEC) with the hippocampus (HP), visual cortex (VC), and auditory cortex (AUC) in fruit bats (left) and baboons (right). In fruit bats: yellow = MEC - HP, red = MEC - AUC, green = HP - AUC, blue = AUC - VC. In baboons: blue = MEC - HP, green = HP - VC, yellow = AUC. **(C)** Normalized and scaled number of tracts between ROIs in both species. **(C)** Visualization of module scores of GO term annotated gene sets on UMAP.

Consistent with the EC’s role as the main cortical input to the hippocampus [7], we identified the angular bundle emerging from the MEC in baboons. In both species, we observed short perforating fibers connecting the MEC to the ventral-anterior hippocampus corresponding to the perforant pathway (Figs. 5A, S7). Together, the angular bundle and the perforant pathway contribute to the trisynaptic hippocampal circuit [8]. Note that in baboons, the fine branching of the angular bundle into hippocampal subregions, (reported in rodents and seen in the fruit bat) could not be resolved, likely due to size limitations when counting tracts between ROIs (Fig. 5C). We also detected anterior projections from the MEC to the perirhinal cortex and the amygdala in baboons (Fig. 5A), consistent with previous reports [25]. In addition, the MECs in two hemispheres in baboons were connected via the hippocampal commissure. In fruit bats, the MEC connectivity also showed additional rostrolateral projections (Fig. 5A), which may have included projections to the LEC. In the baboon, we found no direct connectivity between the visual and auditory primary domains to the MECs (Fig. 5C). Additionally, there was rather weak connectivity between the visual cortex and the hippocampus. While such connections have been suggested in humans [26,27], the signal may also represent connections with the thalamus in our dataset.

In fruit bats, while we also did not observe connections of the visual cortex to the MEC, the auditory cortex was strongly connected with the MEC, hippocampus, and visual cortex, a pattern similar to that reported in rodents [28,29]. Additionally, there was lack of connectivity between the visual cortex and the hippocampus. Thus, the fruit bat MEC exhibits connectivity features more similar to rodents than to primates.

Together, these connectivity patterns reveal species-specific organization of the spatial navigation network that mirrors differences in sensory integration strategies. To understand the cellular and molecular basis of these differences, we next examine neuronal specialization and gene expression programs across species.

### Neuronal specialization and molecular mechanisms underlying conserved and species-specific connectivity patterns in mammalian spatial navigation

To investigate molecular profiles underlying conserved and species-specific connectivity patterns in the spatial navigation system, we assessed enrichment of Gene Ontology (GO) terms related to sensory and cognitive functions across neuronal subtypes in integrated snRNA-seq for primates, mice and fruit bats. Since spatial navigation includes visual, auditory and olfactory cue, we included GO terms related to visual learning, auditory behavior, and olfactory learning, as well as broader learning and memory GO terms (Fig. 6). To further integrate DTI-based large-scale connectivity with molecular profiles, we utilized CellChat [30] that allows prediction of cell-cell connectivity within snRNA-seq data (Fig. 6).

**Figure 6.**
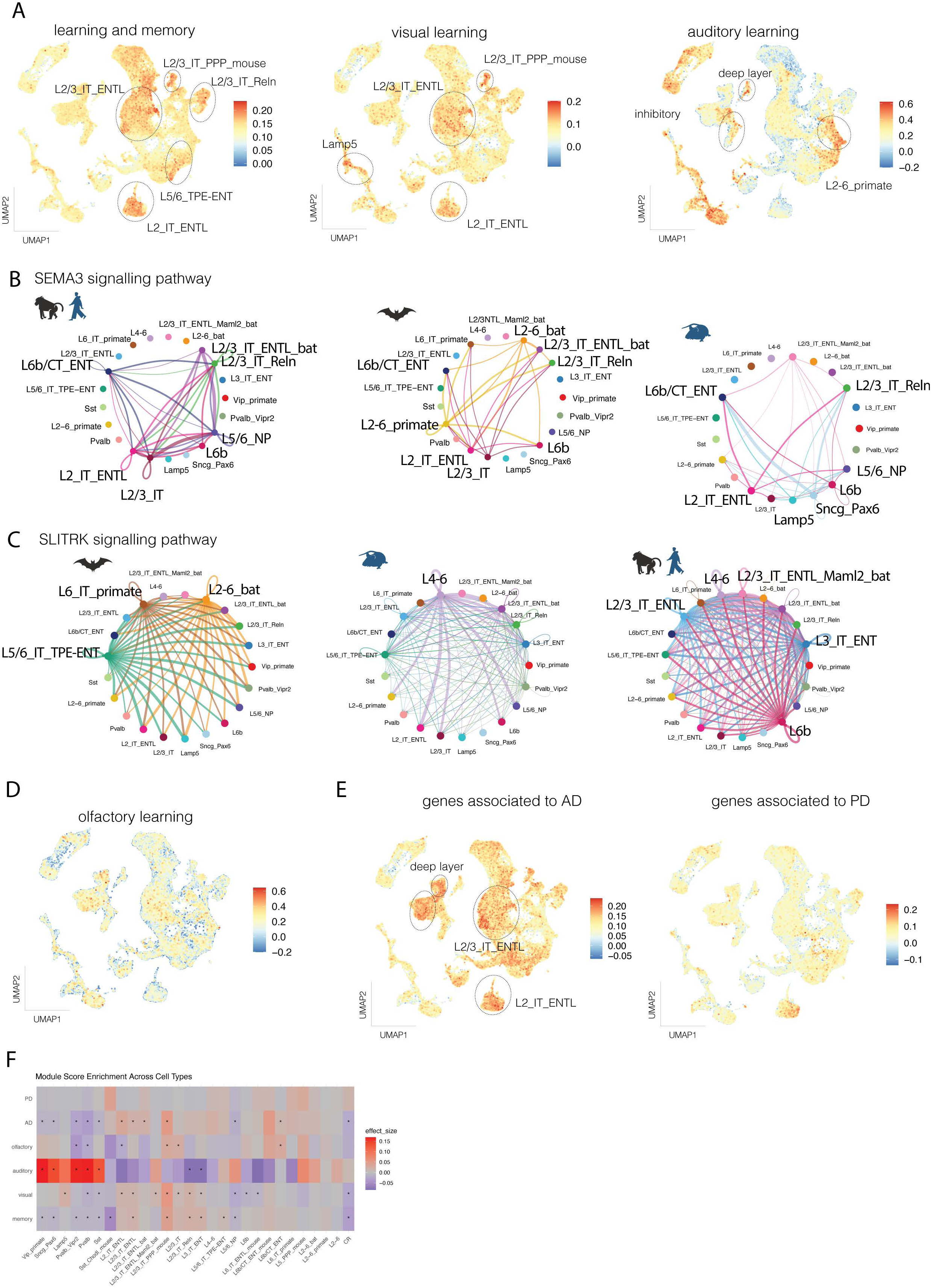
Superficial-layer excitatory neurons and long-range projections link the entorhinal cortex to sensory and cognitive circuits. **(A)** Visualization of module scores of GO term annotated gene sets on UMAP. **(B)** Cell-cell communication of the SEMA3 pathway visualized on circle plots within each species in the integrated dataset. **(C)** Cell-cell communication of the SLITRK pathway visualized on circle plots within each species in the integrated dataset. **(D)** Visualization of module scores on a UMAP of the gene set associated with olfactory learning GO term. **(E)** Module scores of gene sets associated with Alzheimer’s disease (AD) and Parkinson’s disease (PD) visualized on UMAP. **(F)** Heatmap showing pseudobulk module score effect sizes (median differences) across neuronal cell types, where each value represents the difference between a given cell type and all others at the sample level. Adjusted p-values (Benjamini–Hochberg) are overlaid to indicate statistical significance (0.1 >) of the observed enrichment.

As the EC provides the main cortical input to the hippocampus, key region for learning and memory, genes associated with learning and memory were enriched in multiple neuronal subtypes (Fig. 6A). The EC processes various sensory inputs and relays them to the hippocampus [7], and this sensory input processing happens mainly in superficial and deeper layers of the EC [31]. Our data shows predominant enrichment of genes associated with learning or memory in both superficial- (L2_IT_ENTL, L2/3_IT_Reln, L2/3_IT_ENTL) and deep-layer subtypes (L5/6_IT_TPE-ENT) (Figs. 6A,F, S9A). Interestingly, similar enrichment in communication between superficial and deep layer neurons was captured in ligand-receptor expression of the Sema3 pathway (Fig. 6B), which plays a role in regulating synapse formation and function [32]. In species-specific datasets, primates showed higher expression of learning and memory-associated genes in Lamp5 inhibitory subtypes in addition to superficial-layer excitatory neurons (Figs. S8B, S9D, S10C). In mice, all superficial layer-annotated subtypes exhibited high enrichment of learning and memory gene expression signature, while in fruit bats, the highest enrichment was in the L4/5_IT and L2/3_IT_ENTlL_Cux2 subtypes (Figs. S8C,D, S9, S10), suggesting some species-specific EC network that coordinates processing and transfer of sensory input to the hippocampus, overlayed with conserved mammalian EC connectivity.

The visual learning gene set showed the strongest enrichment in superficial-layer subtypes in the cross-species dataset, specifically L2/3_IT_ENTL, L2_IT_ENTL, and L2/3_IT_PPP_mouse (Figs. 6A,F, S9A), mirroring patterns observed in the mouse only dataset (Figs. S8, S9C, S10B). This is consistent with the fact that superficial EC layers receive input from multiple sensory and association cortices [7]. Interestingly, the L2/3_IT_ENTL_Maml2 superficial neurons exhibited enrichment in the bat dataset as well (Figs. S8, S9B, S10A), further supporting specialized spatial information processing by superficial layers. Among inhibitory neurons, the Lamp5 cluster showed strong enrichment in both the cross-species (Fig. 6A,F) and primate-specific (Figs. S8B, S9D, S10C) datasets.

The genes associated with auditory behavior showed strikingly opposite enrichment patterns to visual learning genes. To this end, auditory behavior genes were more strongly enriched in deep-layer excitatory subtypes and in the mixed-layer L2–6_primate subtype, as well as across all inhibitory subtypes in the cross-species integration (Figs. 6A,F, S9A). These trends were consistent in species-specific datasets, including the primate L5/6_IT_TPE-ENT_Cux2 subtype, which substantially contributed to the L2–6_primate cluster (Figs. S8A,B, S9, S10). Surprisingly low module scores for auditory behavior–associated genes in the fruit bat dataset (Figs. S8D, S9B, S10A) may be attributed to the lack of fruit bat genes in existing GO term databases. Mechanistically, deep-layer subtypes across all species were interconnected by the SLITRK signaling pathway (Fig. 6C), which is implicated in synapse formation [33].

The only direct sensory input to the EC in primates that processes a single sensory modality (smell) comes from the olfactory areas [34], and the two are strongly connected in rodents as well [7]. Therefore, we assessed the expression of genes linked to olfactory learning GO term and found no consistent high enrichment in any neuronal subtype across species-specific or cross-species datasets (Figs. 6D,F, S9, S10).

Finally, to show translational potential of our resource, we analyzed enriched gene expression for two neurodegenerative diseases: Alzheimer’s disease (AD), given its relevance to the EC [35,36], and Parkinson’s disease (PD) as non-relevant brain disorder to the EC (Figs. 6E,F, S8E-H, S9, S10). Genes linked to AD were strongly enriched in the EC, in contrast to much weaker enrichment of PD-associated genes, consistent with the well-established involvement of the EC in AD pathology. Interestingly, superficial layer subtypes that were specific to the EC showed the strongest enrichment of AD-associated genes, particularly the L2 _IT_ENTL subtype, in line with its role as a primary site of early tau accumulation and neurodegeneration [37]. The Sema3 pathway, also enriched in L2_IT_ENTL (Fig. 6B), has been explored in therapeutic targeting in AD-related mechanisms [38]. In the primate-specific dataset, the Sst_Chodl inhibitory subtype also showed enrichment for AD-associated genes (Fig. 6E).

Together with the cellular taxonomy, gene set enrichment analyses enhance the functional interpretability of the atlas. By linking conserved and species-specific neuronal subtypes to behavioral, cognitive, and disease-related gene programs, the mammalian EC atlas provides a valuable framework for studying the molecular basis of memory, navigation, and vulnerability across mammals. Finally, user friendly tools for atlas exploration (links will be added in the accepted version) broaden usability and share of the atlas-derived data.

## Discussion

There is a fast-growing number of single-cell transcriptomic atlases generated from various tissues of model organisms [17,39,40]. However, it is challenging to study non-model organisms and even more challenging to cross-compare model and non-model organisms. Through our cross-species EC atlas, we demonstrate that species lacking well-annotated reference genomes or even those that are not included in ortholog databases can still be incorporated into comparative single-cell studies together with model organisms. The use of non-model species is crucial for studies of complex brain functions, since often model organisms (such as laboratory mice) are contained in environments that constrain behavioral complexity. Further, comparative cross-species single-cell transcriptomic studies spanning various evolutionary distances could uncover conserved evolutionary patterns and species-specific innovations [12,14,41]. Here, we study the EC, which is crucial for spatial navigation in mammals. Although previous studies show that EC is a well-conserved brain structure at both the connectivity [7,42,43] and electrophysiological [2] levels across studied mammals, mammalian species differ dramatically in their spatial navigation capabilities. These distinctions are therefore likely to arise from molecular profiles of neuronal subtypes.

Superficial layers of the EC, particularly layers II and III, play a central role in spatial navigation, memory encoding, and associative learning. Projections from superficial layers of the LEC and MEC target the hippocampus and predominantly originate from *Reln+* neurons. These neurons likely correspond to stellate cells in the MEC and fan cells in the LEC [44,45]. Stellate cells are proposed to correspond to grid cells in the MEC, the key spatial navigation neurons [7,46–48]. Grid cells provide spatial information to hippocampal place cells, thus forming a central component of the EC–hippocampus navigational circuit [1,49]. Fan cells on the other hand, contribute to object-locations learning [50], and the processing of object-place-context associations [48] among others. We identified two *Reln+* excitatory subtypes in the cross-species dataset - L/3_IT_Reln likely includes stellate cells, and L2_IT_ENTL includes fan cells (Fig. 2B). While *Reln+* cells in mice and fruit bats were mainly identified in superficial layers (L2_IT_ENTL and L2_IT_ENTm for mouse; L2_IT_ENTL and L2/3_IT_ENTm for fruit bat), primates showed additional *Reln+* expression in deeper layer L5/6_IT_TPE-ENT_Cux2 subtype (Fig. 3B). Interestingly, this expression pattern is not unique to primates as previous reports identified *Reln+* cells in the layer V of the EC of rats and ferrets [31]. Overall, our cross-species atlas captures the key cell types involved in spatial navigation within the EC, linking anatomical identities with transcriptomic profiles and providing insight for spatial navigation from non-model animals.

Bats are the only true flying mammals [6], and during foraging, they cover great distances daily, navigating through diverse terrains [2]. Additionally, fruit bats (*Megachiroptera*) possess an enlarged telencephalon, particularly the neocortex, a similar evolutionary trend observed in primates [46]. In recent years, they have become a model for spatial navigation studies in both laboratory and natural settings [2,10,51–53], although genetic data for this group remain limited. We present the first single-cell dataset from the fruit bat EC, providing a molecular and transcriptomic anchor for existing behavioral, anatomical, and electrophysiological findings. Despite exhibiting several primate-like brain characteristics, and cognitive skills such as cognitive map-based navigation [51], the layer II morphology of fruit bats has been reported to be more rodent-like [46]. Our findings support this observation, with *Reln* expression patterns (and thus potential grid cells) resembling those in mice. Importantly, we also identified several neuronal populations in the superficial layers dominated by fruit bat neurons, which might underlie 3D navigation. Since bats are the only mammals that fly and navigate in 3D space, such demands may drive transcriptomic divergence to neuron types responsible for true 3D navigation. This notion is supported by differences in the properties 3D grid cells recorded in flying fruit bats compared to rodents [10].

Visual cues play a dominant role in primate navigation. Notably, fruit bats possess a primate-like retinotectal pathway, which differs in its organization from that of other mammals, including rodents. Furthermore, a middle temporal visual area has been identified only in fruit bats and primates [54]. This difference was not reflected in our transcriptomic findings. Genes associated with visual learning GO term showed strong expression specifically in superficial layer clusters both in cross-species and species-specific datasets in all species, consistent with visual input primarily targeting layer II and III in mammals [25,55]. In primates, visual sensory input is polymodal [34], which is reflected by the absence of direct connectivity between the MEC and the visual cortex found in baboons in our DTI analysis (Fig 6B,C). While superficial layers receive visual information, layer V is predicted to integrate EC input [7]. It is supported by our data showing that genes linked to learning and memory GO term are expressed not only in superficial layer clusters but also in L5/6_IT_TPE-ENT clusters together with receptor-ligand cell communication of the Sema3 pathway enriched in both superficial and deep layers.

Auditory information provides additional cues for navigation for fruit bats, since for instance Egyptian fruit bats employ echolocation for navigation [56] even in daylight [57]. Our DTI dataset supports the importance of auditory input in navigation as demonstrated by the direct connectivity observed from the auditory cortex to both the MEC and hippocampus in fruit bats. Moreover, we found strong connectivity between the visual and auditory cortices, indicating integration of these sensory inputs in navigation. Since primates are also known to integrate auditory and visual input [34], our results underscore how both fruit bats and primates integrate visual and auditory cues to support spatial navigation. Interestingly, since gene expression signatures, corresponding to visual and auditory information processing and cell-cell communication in the SLITRK signalling pathway, were different between primates and bats in our data, this might underlie species-specific connectivity patterns.

Inhibitory neuronal subtypes were well conserved across all species, with MGE-derived Pvalb and Sst subtypes showing also conservation of relative proportions (Figs. 4C, S4D). Both subtypes modulate principal neurons in the EC [58–60], therefore playing a critical role in spatial representation. Although most studies on these subtypes have been conducted in mice, the distribution of *Pvalb+* cells in the EC appears similar across mammals, including fruit bats [46]. Conversely, CGE-derived inhibitory subtypes exhibited greater divergence, and notably, primates displayed a higher relative abundance of these neurons, with the Vip subtype largely comprising them. CGE-derived Vip inhibitory neurons have been shown to play a specialized role in spatial memory within the MEC by innervating Pvalb and Sst neurons [61] and providing long-range input via the thalamocortical pathway [62]. While these findings are based on studies in mice, the expansion of Vip neurons in primates may underlie the evolution of more complex navigational abilities. This is consistent with the broader expansion of CGE-derived inhibitory neurons across associative cortical areas in humans, a pattern proposed to support increased cognitive capacity [20]. Our EC atlas also captured the expansion of CGE-derived neurons in primates. Strikingly, Vip neurons were nearly absent in our fruit bat dataset, indicating that fruit bats may utilize alternative cellular mechanisms to achieve advanced navigational abilities.

Neurons expressing entorhinal superficial layer (layers II and III) markers were the most abundant in our cross-species dataset. These layers receive diverse cortical inputs, including sensory signals, and project broadly to all subdivisions of the hippocampus [7,34]. The principal neurons described in these layers include stellate (MEC-specific), fan (LEC-specific), and pyramidal cells [7]. The cellular composition of the fruit bat EC is reported to differ notably, with stellate cells (although still being the dominant type) being less abundant. Pyramidal cell abundance in fruit bats was comparable to that in rats and primates, while a surprisingly large population of cells appeared to be neither stellate nor pyramidal [46]. Our transcriptomic data support such cellular composition, where two fruit bat–dominant superficial subtypes (L2/3_IT_ENTL_bat, L2/3_IT_ENTL_Maml2_bat) might reflect this unique transcriptomic specialization of superficial layers in fruit bats. Notably, the *Maml2+* bat L2/3 subtype showed weak expression of canonical superficial EC markers and formed a tightly clustered group distinct from other species in the integrated dataset (Figs. 2, 3B). These fruit-bat specific molecular characteristics of superficial layers potentially illustrate evolutionary innovations in their spatial processing center, which again can be explained by their unique 3D navigation among mammals.

In primates, we also observed distinct gene expression profiles across excitatory layers. Although structural similarities in layer V have been reported between fruit bats and primates [46], the L5/6_IT_TPE-ENT_Cux2 subtype identified in primates expressed *Cux2* (Fig. S2A), a marker typically restricted to superficial layers in rodents. This resulted in a mixed-layer subtype in the cross-species dataset, where superficial and L5/6_IT_TPE-ENT_Cux2 labelled neurons co-clustered (Figs. 3C-D, S4C). This subtype also expressed *Foxp2*, which has been shown to have primate-specific expression pattern in L3-5 IT excitatory neurons [12]. *Cux2* have been reported in mid-to-deep cortical layers in humans and in other primates [12,63]. This highlights that while most excitatory neurons clustered by their predicted cortical layer origin across species, species-specific gene expression patterns influenced conserved layer markers, leading to mixed-layer clusters.

Comparative single-cell analyses facilitate the discovery of evolutionary mechanisms underlying cellular diversity and function. We present a single-cell atlas of the mammalian EC spanning two primates, one rodent, and a fruit bat species, in which we characterized conserved and species-enriched neuronal subtypes. By leveraging cell type annotations from well-characterized model animals, we identified species-specific cell type compositions and gene expression patterns, highlighting how evolutionary pressures such as navigational demands and cognitive specialization have shaped both conserved and lineage-specific transcriptional programs across cortical layers of the EC. Through this atlas, we demonstrate that species without well-annotated reference genomes can be successfully integrated into cross-species analyses. To further contextualize our transcriptomic findings, we explored the connectivity of the MEC in baboons and fruit bats, revealing both shared and divergent features of the spatial navigation system, which we linked to specific neuron types and mechanism underlying these features. Our cross-species atlas offers a valuable resource for investigating the molecular foundations of navigation and memory in mammals. Entorhinal-specific superficial excitatory neurons, highly vulnerable in Alzheimer’s disease [64], consistently enriched for AD-associated genes across species, making this atlas a key reference for studying conserved molecular features across species and their relevance to Alzheimer’s disease.

## Acknowledgements

We are grateful to the members of the Group of Brain Development and Disease Brain and the members of the Khodosevich and Hemberg groups for their critical input. This project was supported by Lundbeckfonden grant (R344-2020-380) to VH and Novo Nordisk Foundation Hallas-Møller Ascending Investigator grant (NNF21OC0067146) to K.K..

## Author contributions

D.M.R., S.E.S., V.H. conceptualized the project. D.M.R. harvested and dissected fruit bat and baboon brain tissue with assistance from S.A.T, M.F.B. and V.H.. D.M.R conducted snRNA-seq experiments with assistance from I.K.. Data analysis was carried out by D.M.R. with assistance from J.J.W and supervision of J.G., M.H., S.E.S., V.H., M.P.W. and K.K.. Immunohistochemistry was carried out by D.M.R. and J.S.. MR imaging and image processing was carried out by Y.M. and D.M.R. with assistance from M.P.W. V.H. and K.K. acquired funding. D.M.R. wrote and revised the manuscript with K.K., M.P.W., J.J.W., M.H., S.E.S, Y.K., S.A.T, M.F.B..

## Declaration of interest

The authors declare no competing interests.

## Methods

### Postmortem baboon and fruit bat brain tissue collection

Tissue samples from Hamadryas baboons (Papio hamadryas) and Egyptian fruit bats (Rousettus aegyptiacus) used for DTI and snRNA-seq were obtained from the Copenhagen Zoo (Zoologisk Have København) (Supplementary Table 1). All animals were clinically healthy and were culled for population management reasons [65].All dissection instruments and Eppendorf tubes were sterilized by autoclave and were sterilized with 70% ethanol and/or RNaze Away between every snRNA-seq sampling.

#### Fruit bat

The whole brain was isolated for DTI and was placed in 4% PFA of formalin. For snRNA-seq, EC was dissected from fresh brains and was snap frozen and stored at -80C. The position of the entorhinal cortex was identified according to the figure from Gatome et al. 2010 [46]. Three incisions were made in a triangle form on the ventral caudal telencephalon by a sterile surgical scalpel and the cortex was lifted carefully off of the underlying white matter and was placed in an Eppendorf tube. Whole brains were kept in 4% PFA for 3 days at 4°C for fixation. Samples were then transferred to long term buffer (0.02% Sodium Azide (Sigma-Aldrich #26628-22-8) in PBS) and were kept at 4°C until imaging.

#### Baboon

For DTI, the brain was perfused through the carotids with 0.5L saline followed by 0.5L 4% PFA for one brain and perfused with 0.5L PBS followed by 0.5L formalin for the last two brains. After perfusion, the skin was removed from the cranium, and the cranium was removed by bone rongeurs chipping away small pieces at the time. The isolated and perfused whole brain was then placed in 4% PFA or in formalin. The position of the EC was identified by gyri on the temporal lobe using the stereotaxic atlas [66] as reference. Two dorsal-ventral incisions were made to produce a coronal brain slice, which subsequently was placed on its posterior surface (facing the anterior surface upwards). The EC was identified on the coronal plane according to the atlas [66]. The entorhinal cortex was then dissected by a sterile surgical scalpel and was placed in a sterile Eppendorf tube. Whole brain perfused with 4% PFA was kept in 4% PFA for 1 week at RT for fixation. The two whole brains perfused with formalin were kept in formalin for 5 days at RT for fixation. Samples were then transferred to long term buffer (0.02% Sodium Azide (Sigma-Aldrich #26628-22-8) in PBS) and were kept at 4°C until imaging.

### NUCLEI ISOLATION

#### Fruit bat

Samples were kept on dry-ice and buffers were pre-chilled on ice. 1mL lysis buffer (10mM TrisHCl, 10mM NaCl (Thermo Scientific #10297760), 3mM MgCl2 (Supelco #105833), 0.01% Tween-20 (Sigma #8221840500), 0.01% NP-40 (Millipore #492016-100), 1% BSA (Thermo Scientific #AM2616), 1mM DTT, 1U/μL RNase inhibitor (Sigma #3335402001), 1x Roche Protease Inhibitor (Roche #11873580001), Nuclease-free water) was added to sample tube and tissue was grinded on ice using a douncer. Next, samples were filtered on 40μm cell strainer (Corning Falcon #CLS352340), tube (Sigma #EP0030108418) and douncer was rinsed with 250μL lysis buffer and that was filtered that as well. Filter was then washed with 250μL lysis buffer. 250μL was transferred from each sample to a new tube (Sigma #EP0030108418) to separate each sample into 2 tubes. 250μL lysis buffer was added to each tube. Samples were then centrifuged at 1,000 x g for 8 min at 4 °C. Centrifuge tubes were coated with 250μL 0.5% BSA. After centrifugation, supernatant was aspirated, and pellet was carefully resuspended in 263μL lysis buffer. Next, 473.4μL of 1.8 Sucrose Cushion Solution (Nuclei PURE 2 M Sucrose Cushion Solution (Sigma #S9308), Nuclei PURE Sucrose Cushion Buffer (Sigma #S9058), 1 M DTT) was added to each resuspended sample and was carefully pipette mixed ten times. BSA was removed from centrifuge tubes and 263μL 1.8 Sucrose Cushion Solution was added to each centrifuge tube. Next, 736.4μL of nuclei sample mixed with sucrose was carefully layered on top of the 263μL 1.8 Sucrose Cushion Solution without mixing. Samples were centrifuged at 13,000 x g for 45 min at 4 °C. Supernatant was aspirated and pellet was resuspended in 50μL of wash/resuspension buffer 10mM TrisHCl, 10mM NaCl (Thermo Scientific #10297760), 3mM MgCl2 (Supelco #105833), 1% BSA, 0.001U/μl RNase inhibitor (Sigma #3335402001), Nuclease-free water). Finally, samples were filtered on 30μm cell strainer (Milteny Biotech #130-098-458) and filter was washed with 20μL wash/resuspension buffer. Nuclei were counted on hematocytometer to calculate concentration.

#### Baboon

Samples were kept on dry-ice and buffers were pre-chilled on ice. 2mL lysis buffer (10mM TrisHCl, 10mM NaCl (Thermo Scientific #10297760), 3mM MgCl2 (Supelco #105833), 0.01% Tween-20 (Sigma #8221840500), 0.01% NP-40 (Millipore #492016-100), 1% BSA (Thermo Scientific #AM2616), 1mM DTT, 1U/μL RNase inhibitor (Sigma #3335402001), 1x Roche Protease Inhibitor (Roche #11873580001), Nuclease-free water) was added to sample tube and tissue was grinded on ice using a douncer. Next, samples were filtered on 40μm cell strainer (Corning Falcon #CLS352340), tube (Sigma #EP0030108418) and douncer was rinsed with 500μL lysis buffer and filtering that as well. Filter was then washed with 500μL lysis buffer. 500μL was transferred from each sample to a new tube (Sigma #EP0030108418) and 500μL lysis buffer was added to each of them. Samples were then centrifuged at 1000 x g for 8 min at 4 °C. Centrifuge tubes were coated with 250μL 0.5% BSA. After centrifugation, supernatant was aspirated, and pellet was carefully resuspended in 263μL lysis buffer. Next, 473.4μL of 1.8 Sucrose Cushion Solution (Nuclei PURE 2 M Sucrose Cushion Solution (Sigma #S9308), Nuclei PURE Sucrose Cushion Buffer (Sigma #S9058), 1 M DTT) was added to each resuspended sample and were carefully pipette mixed ten times. BSA was removed from centrifuge tubes and 263μL 1.8 Sucrose Cushion Solution was added to each centrifuge tube. Next, 736.4μL of nuclei sample mixed with sucrose was carefully layered on top of the 263μL 1.8 Sucrose Cushion Solution without mixing. Samples were then centrifuged at 13,000 x g for 45 min at 4 °C. Supernatant was aspired and pellet was resuspended in 90μL of wash/resuspension buffer (10mM TrisHCl, 10mM NaCl, 3mM MgCl2, 1% BSA, 0.001U/μl RNase inhibitor, Nuclease-free water). Finally, samples were filtered on 30μm cell strainer and filter was washed with 20μL wash/resuspension buffer. Nuclei were counted on hematocytometer to calculate concentration.

### snRNA-seq library preparation and sequencing

Nuclei suspensions were immediately processed on 10x Genomics platform following the manufacturer’s protocol (CG000315 Rev C). 16,000 nuclei were loaded to target 10,000 nuclei recovery on Chromium Controller (10x Genomics). cDNA library preparation was carried out using Chromium Next GEM Single Cell 3’ Kit v3.1 (10x Genomics, PN-1000268) and Dual Index Kit TT Set A (10x Genomics, PN-1000215). Over 20,000 raw reads per nucleus were targeted for sequencing to reach a sufficient sequencing depth. Libraries were pooled and were sequenced on a single lane of Illumina NovaSeq. Samples were demultiplexed after sequencing.

### Accessing published data sets

Raw FASTQ files from published mouse (SRP341866) [17] and human (SRP343046) [14] EC snRNA-seq data were accessed from NIH’s Sequence Read Archive (SRA) (Supplementary Table 2).

### Preprocessing of genome and gene annotation of Egyptian fruit bat

At the time of data analysis, a reference genome assembly for the Egyptian fruit bat (*R. aegyptiacus*) was lacking. As a result, the reference genome Pvam_2.0 (GCF_000151845.1) of the most closely related bat species, the large flying fox (*Pteropus vampyrus*) was used. These two species of flying foxes both belong to the *Pteropodidae* family. However, reference FASTA file of the large flying fox did not include mitochondrial sequences. Therefore, mitochondrial FASTA file was downloaded from https://www.ncbi.nlm.nih.gov/nuccore/NC_026542.1?report=fasta. The mitochondrial FASTA files were reformatted to match the line length of the (genomic) reference FASTA file. Finally, the two FASTA files were concatenated.

The reference GTF file of the large flying fox also lacked mitochondrial gene annotations. Mitochondrial annotation was therefore manually generated for the 13 protein-coding genes. The information was extracted from https://www.ncbi.nlm.nih.gov/nuccore/NC_026542.1?report=genbank and the baboon GTF line format was used to produce the mitochondrial GTF file for the fruit bat.

### Preprocessing of genome and gene annotation of baboon

Similar to the Egyptian fruit bat, the hamadryas baboon (*P. hamadryas*) also lacked a reference genome assembly at the time of data analysis. Therefore, the reference genome Panubis1.0 (GCF_008728515.1) of the closely related olive baboon (*P. anubis*) was used. Before gene expression quantification by Cell Ranger, a custom reference has to be built when no pre-built reference is available. During this process, GTF files are filtered to retain only genes of interest (i.e., protein-coding genes). This information is found in the attribute column of GTF files. This column contains a list of tag-value pairs. Whether a gene is protein-coding is indicated by the biotype tag. Cell Ranger’s mkgtf filter function requires the biotype tag to appear in the same position within the attribute column for every gene entry in the GTF file. However, the reference GTF files we used did not have the biotype tag in a consistent position and therefore they required filtering with a custom parser prior to custom reference building with Cell Ranger. In brief, gene IDs corresponding to protein-coding genes were extracted from the GTF files and compiled into lists, and GTF files were filtered based on these gene lists. The resulting GTF files were formatted correctly for subsequent steps.

### Single nuclei expression quantification and data preprocessing

Cell Ranger (version 7.1.0) *mkref* command was run on the corrected and filtered reference genomes producing custom built references for baboons and fruit bats for subsequent read alignment and counting. Following the standard Cell Ranger workflow, reads from each sample were aligned to their respective references and gene expression was counted by running Cell Ranger’s *count* command. We incorporated published single-nucleus expression data from human [14] and mouse [17] entorhinal cortex. These published datasets were generated using the same platform as the one used for our samples (10x Genomics). In case of published data, the available pre-built references (refdata-gex-mm10-2020-A, refdata-gex-GRCh38-2020-A, respectively) provided by 10x Genomics were used in the mapping and counting step. The Cell Ranger web summary was used to assess mapping statistics and read quality. One of the fruit bat samples did not pass our quality filters and was not used in further steps. The remaining filtered raw count matrices were used for subsequent analyses. These matrices were then imported in R (version 4.4.1) and were transformed into Seurat objects (Seurat version 5). “Raw” count matrixes were saved at this point as Seurat objects for ortholog mapping. First, low quality cells were filtered out with a higher than 5 percent of mitochondrial gene fraction and higher than 2 for fruit bats, or too high or low gene and UMI counts (Figure S9A). Second, doublets were removed via DoubletFinder (version 2.0.4) [67]. Third, each biological replicate was normalized by SCTransform package (version 0.4.1), and UMAPs were inspected (Fig. S9B). Concatenated Seurat objects for each species were saved for ortholog mapping. One of the baboon samples did not contain enough high-quality cells to be able to continue further analysis and was discarded. Biological replicates were then integrated using the RPCA method in Seurat. At this step, each species had three biological replicates. Batch effects on clusters were visually inspected on UMAPs after integration (Fig. S10A-D).

### Ortholog mapping

Gene orthologs across our target species were mapped by OrthoFinder [68]. For that, the proteomes were downloaded from Ensembl for the following species: mouse, rat, zebra fish, human, bonobo, chimpanzee, gorilla, macaque, baboon, fruit bat, salmon. Following the standard OrthoFinder workflow (https://davidemms.github.io/orthofinder_tutorials/running-an-example-orthofinder-analysis.html), proteomes were filtered. Mitochondrial protein sequences from salmon’s ICSASG_v2 assembly (GCF_000233375.1) were attached to the filtered salmon genomic proteome. For fruit bat, protein sequences of mitochondrial genes were downloaded from the same mitochondrial assembly (https://www.ncbi.nlm.nih.gov/nuccore/NC_026542.1) used for reference genome curation and were attached to the filtered fruit bat genomic proteome. Gene duplicates were filtered out from human, mouse and zebrafish filtered proteomes. To map orthologs between human and baboon, OrthoFinder (version 2.5.5) was run on the seven primate species, and to map orthologs across our four mammalian target species, additional rodent (rat) and primate (bonobo) species were included along with two non-mammalian vertebrates (zebra fish and salmon). One-to-one orthologs were extracted from OrthoFinder output between: human and baboon, human and mouse, baboon and mouse, fruit bat and mouse. Then, one-to-one gene orthologs were filtered to only contain genes that were found in our “raw” Seurat object count matrices. Non-one-to-one orthologs (many-to-many, many-to-one, one-to-many) were converted to one-to-one orthologs. Orthologs with one gene per species present in our “raw” Seurat object count matrices were regarded as one-to-one. Genes with the highest SCTransform normalized average expression were selected when multiple genes were present in a species from the same ortholog group. For that, expression values were extracted from our preprocessed data (concatenated Seurat objects mentioned in the previous section). Baboon and fruit bat count matrices were imported to R again and the newly created Seurat objects were translated to human and mouse gene orthologs, respectively. These translated Seurat objects were then preprocessed in the same manner as before. For subsequent analysis, these translated preprocessed Seurat objects were used for baboons and fruit bats.

### Major cell type annotation

Major cell types were annotated using canonical marker genes in each species. For baboons and humans, we identified seven major cell types: excitatory neurons (SLC17A7+), inhibitory neurons (GAD1+), astrocytes (AQP4+, APOE+), oligodendrocytes (OLIG1+, OLIG2+, PLP1+, OPALIN+), oligodendrocyte progenitor cells (PDGFRA+), microglia (C1QB+, CX3CR1+), and endothelial (VWF+, FLT1+, MECOM+, CLDN5+, ABCB1+). For fruit bat, we identified five major cell types: excitatory neurons (Slc17a7+), inhibitory neurons (Gad1+, Gad2+, Sst+), oligodendrocytes (Mbp+, Plp1+, Opalin+), oligodendrocyte progenitor cells (Pdgfra+) and closely clustered VLMC and astrocytes (Apoe+, Aqp4+, Ptgds+, Gja1+, Selenop+, Sparc+, Slc15a2+). Neuronal cell types were then subset from each species for neuronal subtype annotation. The published mouse data set only consisted of neuronal cells. The neuronal subsets were stripped using DietSeurat to retain only raw counts and were then renormalized by SCTransform [69] and visualized again on UMAP. However, primate (humans and baboons) data sets were integrated by RPCA method after normalization (S10E).

### Constructing EC reference data set

To facilitate neuronal subtype annotation, we constructed an EC reference dataset by subsetting the Allen Brain Institute’s Mouse Whole Cortex and Hippocampus 10x Genomics transcriptomic dataset [17] to include only cells from the EC. This EC-specific dataset was loaded into Seurat, and a neuronal subset was created. The resulting neuronal subset was normalized and clustered using the same parameters as our experimental samples. Label transfer from the EC reference dataset to each neuronal subset was performed using Seurat’s FindTransferAnchors, TransferData, and AddMetaData functions implementing default parameters.

### Neuronal subtype annotation within and across species

The fruit bat neuronal subset was renormalized and clustered. Human and baboon neuronal subsets were converted to mouse gene orthologs before renormalization and were integrated before clustering resulting in a primate neuronal data set. As mentioned before, neuronal subtypes were mapped to the reference EC data set for each species neuronal subset (mouse, fruit bat, and primate). For each cluster, this mapped cell identity together with marker gene expression were used to annotate neuronal subtypes.

Inhibitory neuronal subtypes were confidently identified by canonical markers. MGE derived neurons expressed the markers *LhxC* and *SoxC*, while neurons originating from CGE were defined by *Adarb2*, *Prox1* and *Nr2f2,* and in fruit bats *Cnr1* expression. We annotated the following inhibitory subtypes in all species: Sst (*Sst+*), Pvalb (*Pvalb+*), Pvalb_Vipr2 (*Pvalb+, Vipr2+*), Lamp (*Lamp5+*), Lamp5_Lhx6 (*Lamp5+, Lhx6+*). In mice and primates, we further identified Sst_Chodl (*Sst+, Chodl+, Npy+*), Vip (*Vip+*) subtypes. Fruit bats had closely clustered *Vip+* and Sncg+ neurons, while we annotated a separate Sncg (*Sncg+*) cluster in mice and its primate homologous cluster (*Kcng1+, Pax6+, Adarb2+*).

For excitatory neurons, upper layers were described by *Cux2* expression, mid to deep layers were *Rorb+* while deep layers expressed *Tle4*, *Bcl11b*, and *Fezf2*. Information about anatomical region (ENT, entorhinal; ENTL, entorhinal lateral; ENTm entorhinal medial, PPP, post-presubiculum parasubiculum; TPE, temporal association-perirhinal-ectorhinal; CTX, isocortex) and axonal projection (IT, intratelencephalic; NP, near-projecting, 6b) for subtypes was added from reference label transfer. We annotated in total 29 unique excitatory neuron subtypes in primates, fruit bats, and mice.

These previously assigned neuronal subtype labels were carried over to cross-species integration, and this information coupled with established marker gene expression was used to determine shared neuronal subtypes.

### Integration across species

Each species consisted of three biological replicates. Cross-species integration was performed using Seurat’s RPCA method (Fig. S10F). Primates (baboons and humans) were integrated for neuronal subtype annotation, as mentioned above. Finally, all annotated mammalian neurons (humans, baboons, mice, and fruit bats) were integrated as well.

### Benchmarking integration methods

To assess how different integration methods affect shared clusters in cross-species integration, we integrated our target species using different methods: Harmony [70], scVI [71] and Seurat’s CCA [72]. For Harmony and CCA, we took the SCTransform normalized samples as input and used the same number of principal components (30) as in RPCA-based integration. Before integrating with scVI, we renormalized samples via *NormalizeData* function, identified top variable genes by *FindVariableFeature*, scaled data through *ScaleData* and ran *RunPCA*. Each integration method was applied using Seurat’s IntegrateLayers function. We integrated both biological replicates and across species by all four integration methods (Fig. S11A). To quantify quality of different integration methods, we calculated metrics using the *getIntegrationMetrics* function from the scIntegrationMetrics package (version 1.2.0) [73] (Fig. S11B).

### Relative proportion of subtypes in each species

Absolute ratios of species within a subtype can be biased by sample size differences; therefore, we calculated normalized proportions of all subtypes in each species. First, we calculated the proportion of subtypes in each sample in the cross-species Seurat object. Then, we took the mean of proportions for each species. Lastly, we also calculated standard deviation for all subtypes in each species.

### Relative contribution from species to subtypes

To correct for the bias that different sample sizes introduce, we calculated the relative contribution of each species to subtypes. We used the previously calculated mean proportions of subtypes within species to compute the overall mean proportion of each subtype for each species. We pooled human and baboon samples to present primates as species for subtype annotation. Finally, when the calculated relative contribution was above 75% in a subtype for a single species, we annotated that subtype as species enriched.

### Subtype annotation within and across species

When subtype annotations were complete in both species and cross-species data sets, neuronal subtype transitions between species-specific and cross-species datasets were visualized using Sankey diagrams. Link data frames were constructed by calculating the number of nuclei in each subtype in each species and in cross-species Seurat objects, then calculated the number of transitions between species subtypes and cross-species subtypes in a pairwise manner for all species. For clearer visualization, transitions with low cell counts (for cortical layers under 50 and for others under 10) were filtered out. To visualize transitions of two species at the same time, we joined two link data frames before generating Sankey diagrams using R package *networkD3*.

### Detection of subtype marker genes

Subtype marker genes were identified using Seurat’s *FindMarkers* function with the default Wilcoxon Rank Sum test in both the cross-species and species-specific datasets. Marker gene detection was performed separately for excitatory and inhibitory subtypes, i.e., gene expression was compared within each neuronal class independently. Marker genes were defined with false discovery rate below 0.01, expression in more than 25% of nuclei in the target subtype, and log2 fold change greater than 1.

### Detection of differentially expressed genes across species within subtypes

In the cross-species dataset, DEGs were identified within each subtype across species. We used the same Seurat function *FindMarkers* function with the default Wilcoxon Rank Sum test. For the final DEGs list, we implemented the following filters: false discovery rate below 0.01, expression in more than 25% of nuclei in the target subtype, and log2 fold change greater than 1. To capture species- and subtype-specific marker genes, we intersected DEG lists from within-species subtype comparisons with DEG lists from cross-species subtype comparisons. In this analysis, primates were treated as a single species. Marker genes were then visualized on heat maps (*DoHeatmap*) and on dot plots (*DotPlot*).

### Expression of GO term annotated gene sets

Gene Ontology (GO) annotations were selected based on their relevance to our research question: navigation and sensory input. We inspected the expression of gene sets annotated to these selected GO terms, namely learning and memory (GO:0007611), visual learning (GO:0008542), auditory behavior (GO:0031223), and olfactory learning (GO:0008355). Gene lists were downloaded from https://www.ebi.ac.uk/QuickGO/annotations. In addition, the EC-relevant Alzheimers’s disease related genes from WP_ALZHEIMERS_DISEASE (https://www.gsea-msigdb.org/gsea/msigdb/mouse/geneset/WP_ALZHEIMERS_DISEASE) and non-relevant Parkinson’s disease associated genes from WP_PARKINSONS_DISEASE mouse gene set (https://www.gsea-msigdb.org/gsea/msigdb/mouse/geneset/WP_PARKINSONS_DISEASE.html) were downloaded. We used *AddModuleScore* in Seurat to assign a score to each cell representing normalized expression rates of our target gene lists. Expression of the target gene lists was scored in the cross-species data set, and the species data sets at a single cell level.

### Transcriptomic divergence of subtypes across species

We calculated and visualized transcriptomic divergence between each pair of species in each subtype. For that, we followed a previously published protocol from Ma et al. [12]. In short, we took the average expression of the top 1500 highly variable genes in each subtype for each species. Then, for each species pair, we calculated transcriptomic divergence as 1 minus the Pearson correlation coefficient. These divergence values were then visualized as a heatmap using *gg_heat* function of *ggplot2* package.

### Cell-cell communication with CellChat

The integrated EC atlas was first sub setted by species (primates, mouse, fruit bat) and only those clusters were kept that had at least 10 nuclei from each species. Next, following the Cellchat vignette, cellchat objects were created from each species subset separately using the package *CellChat* [30]. Then we compared the datasets pairwise following the “Comparison_analysis_of_multiple_datasets” vignette provided by CellChat. Pathways were visualized on circle plots by *netVisual_aggregate*.

### Vibratome sectioning and immunohistochemistry

Collected fixed brains were sliced using a vibrating microtome (Leica VT1000S). Brains were sliced free-floating with 50 microns thick coronal sections and were stored in PBS with sodium azide at 4℃. Selected sections were washed in PBS and permeabilized/blocked for 2 hours at room temperature in PBS containing 0.2% Triton X-100 (VWR 131 International, Denmark) and 10% normal donkey serum (Gibco 16050-122). Sections were incubated overnight at 4°C with primary antibody against sheep anti-VGLUT1 (1:100, Abcam ab79774), goat anti-FOXP2 (1:100, Novus NB100-55411), mouse anti-CUTL2 (1:100, Novus NBP3-26842), mouse anti-calretinin (1:2000, Swant 6B3), rabbit anti-GPD1 (1:200, Proteintech13451-1-AP) in PBS containing 0.2% Triton X-100 and 2% serum. After washing, sections were incubated overnight at 4°C with secondary antibody (1:1000, Invitrogen and Jackson) in the same buffer. Nuclei were counterstained with DAPI (1:50000, Sigma) for 10 minutes at room temperature. Sections were washed and mounted on Superfrost microscopic slides using FluoroMount-G medium (ThermoFischer Scientific) or Immu-mount (Epredia). Mounted sections were stored in a 4℃ fridge in a closed container to protect them from light.

### Imaging

Imaging was carried out using Zeiss confocal microscope LSM800. The 25x objective with 0.5x magnification was used for all images. Z-axis was set for maximum intensity of signals. Consistent parameters were maintained across all images. All Images were analysed in Fiji [74].

### MR imaging

All MRI scanning was performed with a 9.4 Tesla preclinical scanner interfaced to a Bruker Avance III console and controlled by ParaVision 6.0.1 software (Bruker BioSpin, Ettlingen, Germany) at Panum Preclinical MRI Core Facility, University of Copenhagen. To get the best MRI signals from the brain, the PFA-fixed brain samples were immersed in phosphate buffered saline (PBS; 0.1 M, pH 7.4) for rehydration more than a week prior to MRI session, and the samples were suspended into plastic containers filled with a proton-free perfluorinated susceptibility-matching fluid (Fluorinert, 3M, USA) prior to imaging to minimize susceptibility artifacts.

### MRI for fruit bat

The combination of a 1500mT/m gradient coil (BFG6S, Bruker) and a 35-mm inner diameter transmit-receive volume coil (Bruker) was used for fruit bat brain scanning. For the structural references with high spatial resolution, the imaging protocol used a 3D constructive interference in steady-state (CISS) with acquisition of four 3D-TrueFISP volumes of orthogonal phase encoding direction, and other imaging parameters were set to: repetition time (TR) = 5.1 ms, echo time (TE) = 2.55 ms, flip angle = 50°, field of view = 32.1 mm × 19.2 mm × 19.2 mm, matrix = 428 × 256 × 256, image resolution = 75 µm isotropic, number of signal acquisition repetitions = 8. For diffusion tensor imaging (DTI), a 2D Stejskal-Tanner sequence (TR/TE = 3000/20 ms, field of view = 19.2 mm × 19.2 mm, matrix = 96 × 96, slice thickness = 0.4 mm, slice numbers = 96, number of signal acquisition repetitions = 4) was used. We applied the motion probing gradients (MPGs) in twenty non-collinear directions with two b-values (b = 800 and 2000 s/mm2) in addition to 1 b0.

### MRI for baboon

The combination of a 100mT/m gradient coil (B-GA20S, Bruker) and a 154-mm inner diameter transmit-receive volume coil (Bruker) was used for baboon brain scanning. For CISS, four 3D- TrueFISP volumes were acquired with: TR/TE = 5.18/2.59 ms, flip angle = 50°, field of view = 90 mm × 76.8 mm × 76.8 mm, matrix = 300 × 256 × 256, image resolution = 300 µm isotropic, number of signal acquisition repetitions = 8. For DTI, a 2D Stejskal-Tanner sequence (TR/TE = 7000/31 ms, field of view = 96 mm × 96 mm, matrix = 192 × 192, slice thickness = 2 mm, slice numbers = 44, number of signal acquisition repetitions = 4) was used. We applied MPGs in twenty non-collinear directions with two b-values (b = 800 and 1600 s/mm2) in addition to 1 b0.

### MR image processing

For CISS, every 3D-TrueFISP volume acquired with 32 repetitions (4 orthogonal phase encodings x 8 image repetitions) was motion-corrected with ANTs [75]. CISS image was computed as an averaged maximum intensity projection from motion-corrected TrueFISP volume, resulting in an image of removed banding artifacts. To correct for B0 inhomogeneities, image bias field correction was conducted using N4 bias field correction algorithm [76]. For DTI imaging, every MPG volume was motion-corrected with ANTs [75]. To suppress the noise in each repetition series, the Marcenko-Pastur PCA (MP-PCA) algorithm denoising has been performed and averaged four repetition series. Diffusion Toolkit (version 0.6.4.1) and TrackVis (version 0.6.1) were used for 3D reconstruction of white matter tracts [77]. We used a DTI mask threshold and an angular threshold of 30 degrees.

### ROI annotation in fruit bats

The Stereotaxic Brain Atlas of the Egyptian Fruit Bat [78] was used as the main reference for ROI annotation. To annotate the MEC and the hippocampus, ROIs were hand-drawn on high resolution T2 weighted images in ITK-SNAP (version 4.2.0) first on the coronal plane followed by fine tuning segmentation on sagittal and horizontal planes. Next, segmentation information was converted to respective b0 images that were acquired for DTI (see data acquisition protocol).

In case of the MEC, the following anatomical structures served as landmarks: rhinal fissure, hippocampus, dentate gyrus, lateral entorhinal cortex, subiculum, pre-, and para-subiculum. First, the rhinal fissure was considered a clear indicator of dorsolateral boundary of LEC, continuing as dorsolateral boundary of the MEC as well. The rostral boundary of the MEC was identified inferior to subiculum at the very caudal part of dentate gyrus. There is no clear boundary between the LEC and the MEC at this resolution, therefore this boundary was determined by noting the disappearing subiculum and simultaneously growing parasubiculum. The MEC was traced to the caudal end of the telencephalon. Lastly, sagittal and horizontal planes were used to fine tune the boundaries of the MEC in relation to the hippocampus and dentate gyrus (Fig. S6A).

To annotate the hippocampus, we followed the clearly visible distinct laminar pattern of this structure. Dorsal, ventral and lateral boundaries were defined by the lateral ventricle or its remnants, the dorsal third ventricle, and caudate respectively (Fig. S6B).

To annotate the visual and auditory cortices, the lower resolution b0 images were used to directly draw ROI’s based on the atlas. The position and extend of the retrosplenial cortex, presubiculum and hippocampus were the major anatomical landmarks used to delineate these sensory cortical areas in the coronal plane (Fig. S6C,D).

### ROI annotation in baboons

The rhesus monkey brain in stereotaxic coordinates [66] was used as the main reference for ROI annotation. To annotate the MEC, ROIs were hand-drawn on high resolution T2 weighted images in ITK-SNAP (version 4.2.0). Similarly to the annotation in the fruit bat, tracing was done first in the coronal plane and finished in sagittal and horizontal planes. Next, segmentation information was converted to respective b0 images. Based on this converted segmentation, final segmentations were drawn on b0 images that were acquired for DTI. The visibility of marked lamination served as the primary marker to identify the MEC. Additionally, the anterior boundary was established with the presence of a well-defined amygdala, and the posterior boundary was marked at the appearance of LGN (Fig. S5A).

The distinct laminar pattern of the hippocampus as well as its border with the lateral ventricle was clearly visible, allowing the delineation of the hippocampus (Fig. S5B).

The auditory cortex, including the primary and secondary domains, was identified using the hippocampus and the putamen as anatomical landmarks, using atlas derived ROI borders (S5D). The visual cortex, also including primary and secondary regions, was identified in the occipital lobe, the appearance of lateral ventricle serving as its rostral boundary (Fig. S5C).

### Analyzing connectivity between ROIs

Diffusion tractography images were loaded into TrackVis (Version 0.6.1), together with previously hand-annotated ROI segmentation data. Connectivity was set to Either end for each ROI, except for the hippocampus in baboons (Any parts), to adjust MEC-hippocampus connectivity detection due to resolution limitations. Tracks between ROIs, as well as those starting or ending in individual ROIs, were counted by the software. These tract counts were normalized by dividing the total number of detected tracts in the given sample and scaled by multiplying by 100. Finally, the mean (replicates n=3) of normalized scaled counts was visualized on bar plots with standard deviation error bars.

## Data and code availability

SnRNA-seq data generated in this study will be deposited at publicly available depositories upon acceptance and the original code, count matrices and Seurat objects are deposited at https://github.com/doreszr/Mammalian_-EC-snRNA-seq-atlas.

## Declaration of generative AI and AI-assisted technologies in the manuscript preparation process

During the preparation of this work the author(s) used ChatGPT in order to improve grammar and style. After using this tool/service, the author(s) reviewed and edited the content as needed and take(s) full responsibility for the content of the published article.

## Notes

https://github.com/doreszr/Mammalian_-EC-snRNA-seq-atlas/tree/main

### Competing Interest Statement

The authors have declared no competing interest.

