## Supplementary figures and text for "Spatial navigation through evolution: a single-cell atlas of the mammalian entorhinal cortex"

Supplementary Data Information

**Supplementary Figures**

**Supplementary Fig. 1 – Major cell types identified in mammalian EC datasets. A)** Major cell types identified in the EC of baboons (left) and fruit bats (right). **(B)** Major cell types identified in the published human (left) and mouse (right) EC snRNA-seq datasets.

**Supplementary Fig. 2 – Expression of selected marker genes visualized on UMAPs. A)** Expression of excitatory layer markers that are uniquely co-expressed in primate neuronal dataset. **(B)** Expression of the same layer markers in the mouse dataset. **(C)** Expression of the same layer markers in the fruit bat dataset. **(D)** Expression fruit bat-specific excitatory and CGE-derived inhibitory marker genes in the fruit bat neuronal dataset. **(E)** Expression of conserved L5/6 NP subtype markers in the cross-species integrated dataset. **(F)** Expression fruit bat-specific excitatory marker genes in the cross-species dataset.

**Supplementary Fig. 3 – Transcriptomic divergence across mammals within conserved entorhinal cortex cell types. (A)** Scaled transcriptional divergence scores across species in the shared neuronal subtypes. **(B)** The top 10 DEGs for each species, was detected in two conserved subtypes: Pvalb (top) and L2/3_IT_ENTL (bottom) (30 DEGs per subtypes). Although these DEGs were captured between species in a specific subtype, their expression remained species-specific across all neuronal subtypes when visualized on dot plots. Neuronal subtypes were subset species-wise to visualize species-specific expression patterns. Subtype order (on the bottom) is consistent across species to enable cross-comparison.

**Supplementary Fig. 4 – Divergence of neuronal cell types across mammals. (A)** Expression of deep layer excitatory subtype markers across species visualized on a heatmap. **(B)** Heat map of differentially expressed genes (DEGs) among mixed-layer subtypes identified in the cross-species integration, illustrating subtype-specific expression patterns. **(C)** Transition of excitatory neurons from mouse-, and primate-specific datasets to cross-species integrated dataset (left). Subtype clusters are grouped according to superficial, deep, and mixed layer identity and CR cells. On the right, origin of neurons in the cross-species mixed L2-6 primate subtype. **(D)** Relative inhibitory subtype proportions in each species. This plot visualizes the relative proportions of subtypes within inhibitory neurons so the proportions in a given species add up to 100%. **(E)** Heat map displaying species-specific differentially expressed genes (DEGs) within conserved inhibitory subtypes.

**Supplementary Fig. 5 – ROI annotation in baboons.** ROIs are shown on the coronal (left) and horizontal (right) planes. **(A)** Medial entorhinal cortex. **(B)** Hippocampus**. (C)** Visual cortex including primary and secondary areas. **(D)** Auditory cortex including primary and secondary areas.

**Supplementary Fig. 6 – ROI annotation in fruit bats.** ROIs are shown on the coronal (left) and horizontal (right) planes. **(A)** Medial entorhinal cortex. **(B)** Hippocampus**. (C)** Visual cortex including primary and secondary areas. **(D)** Auditory cortex including primary and secondary areas.

**Supplementary Fig. 7 – MEC connectivity of biological replicates. (A)** DTI tractography of the MEC connectivity in the other two baboon biological replicates. **(B)** DTI tractography of the MEC connectivity in the other two fruit bat biological replicates. Anatomical planes: R - rostral, C - caudal, S - superior, I - inferior.

**Supplementary Fig. 8 – Module scores of selected gene sets visualized on UMAPs.** Module scores of gene sets linked to GO terms (on the top of the UMAP plots) in the cross-species dataset **(A),** primate dataset **(B),** mouse dataset **(C)** and fruit bat dataset **(D)**. Module scores of gene lists linked to Alzheimer's (AD) and Parkinson's disease (PD) in the cross-species dataset **(E),** primate dataset **(F),** mouse dataset **(G)** and fruit bat dataset **(H)**.

**Supplementary Fig. 9 – Module scores of selected gene sets visualized on violin plots.** Module scores of gene sets linked to GO terms and diseases (on the top of the violin plots) in the cross-species dataset **(A),** fruit bat dataset **(B),** mouse dataset **(C)** and primate dataset **(D)**.

**Supplementary Fig. 10 – Effect sizes of module scores in the species-specific data sets visualized on a heatmap.** Heatmap showing pseudobulk module score effect sizes (median differences) across neuronal cell types, where each value represents the difference between a given cell type and all others at the sample level. Adjusted p-values (Benjamini–Hochberg) are overlaid to indicate statistical significance (0.1 >) of the observed enrichment. **(A)** fruit bat dataset, **(B)** mouse dataset, **(C)** and primate dataset.

**Supplementary Fig. 11 – Preprocessing baboon and fruit bat datasets. (A)** Violin plot of read count, gene count and UMI count of baboon and fruit bat samples before quality filtering. **(B)** UMAP visualization of baboon and fruit bat samples after quality and doublet filtering.

**Supplementary Fig. 12 –Integration of samples after neuronal sub setting, visualized on UMAPs. (A-D)** Integration of biological replicates after neuronal sub setting in the species-specific datasets. **(E)** Cross-species integration of neuronal cells colored by biological replicates.

**Supplementary Fig. 13 – Different cross-species integration methods.** Four integration methods were tested on the primate dataset including glial and neuronal cells. The four methods were Seurat's RPCA and CCA, Harmony, scVI. **(A)** Different integration methods visualized on UMAPs**. (B)** Comparing the four methods by metrics. PCA stands for unintegrated PCA embeddings. The metrics are: iLISI - Local Inverse Simpson's Index (LISI), CiLISI - cell-type aware version of iLISI, cLISI - LISI is calculated on the cell type labels, celltype_ASW - Average Silhouette Width by celltype [73].

**Supplementary Tables**

**Supplementary Table 1 – Metadata of collected samples.**

**Supplementary Table 2 - Sample metadata of published mouse and human data sets.**

Supplementary Table 1 – Metadata of collected samples.

| Sample ID | Age (years) | Collected | Sex |
|---|---|---|---|
| baboon1 | 2.5 | EC left | Female |
| baboon2 | 2 | EC left | Female |
| baboon3 | 1.5 | EC right | Female |
| baboon4 | 3.5 | Whole brain | Female |
| baboon5 | 1.5 | EC right | Female |
| baboon6 | 1.5 | Whole brain | Female |
| baboon7 | 3 | Whole brain | Female |
| bat2 | 1-2 | EC left | Female |
| bat3 | 1-2 | EC left | Female |
| bat4 | 1-2 | EC right | Female |
| bat5 | 1-2 | EC right | Female |
| bat6 | 1-2 | Whole brain | Female |
| bat7 | 1-2 | Whole brain | Female |
| bat8 | 1-2 | Whole brain | Female |

Supplementary Table 2 - Sample metadata of published mouse and human data sets.

| SRA ID | Sample ID | Link |
|---|---|---|
| SRR16928876 | human1 | [73] |
| SRR16928884 | human2 | https://trace.ncbi.nlm.nih.gov/Traces/?view=run_browser&acc=SRR16928884&display=data-access |
| SRR16928886 | human3 | https://trace.ncbi.nlm.nih.gov/Traces/?view=run_browser&acc=SRR16928886&display=data-access |
| SRR16409869, SRR16409878 | mouse1 | https://trace.ncbi.nlm.nih.gov/Traces/index.html?view=run_browser&acc=SRR16409869&display=data-access<br>https://trace.ncbi.nlm.nih.gov/Traces/?view=run_browser&acc=SRR16409878&display=data-access |
| SRR16409870, SRR16409880 | mouse2 | https://trace.ncbi.nlm.nih.gov/Traces/index.html?view=run_browser&acc=SRR16409870&display=data-access<br>https://trace.ncbi.nlm.nih.gov/Traces/index.html?view=run_browser&acc=SRR16409880&display=data-access |
| SRR16409871, SRR16409881 | mouse3 | https://trace.ncbi.nlm.nih.gov/Traces/index.html?view=run_browser&acc=SRR16409871&display=data-access<br>https://trace.ncbi.nlm.nih.gov/Traces/index.html?view=run_browser&acc=SRR16409881&display=data-access |

S1

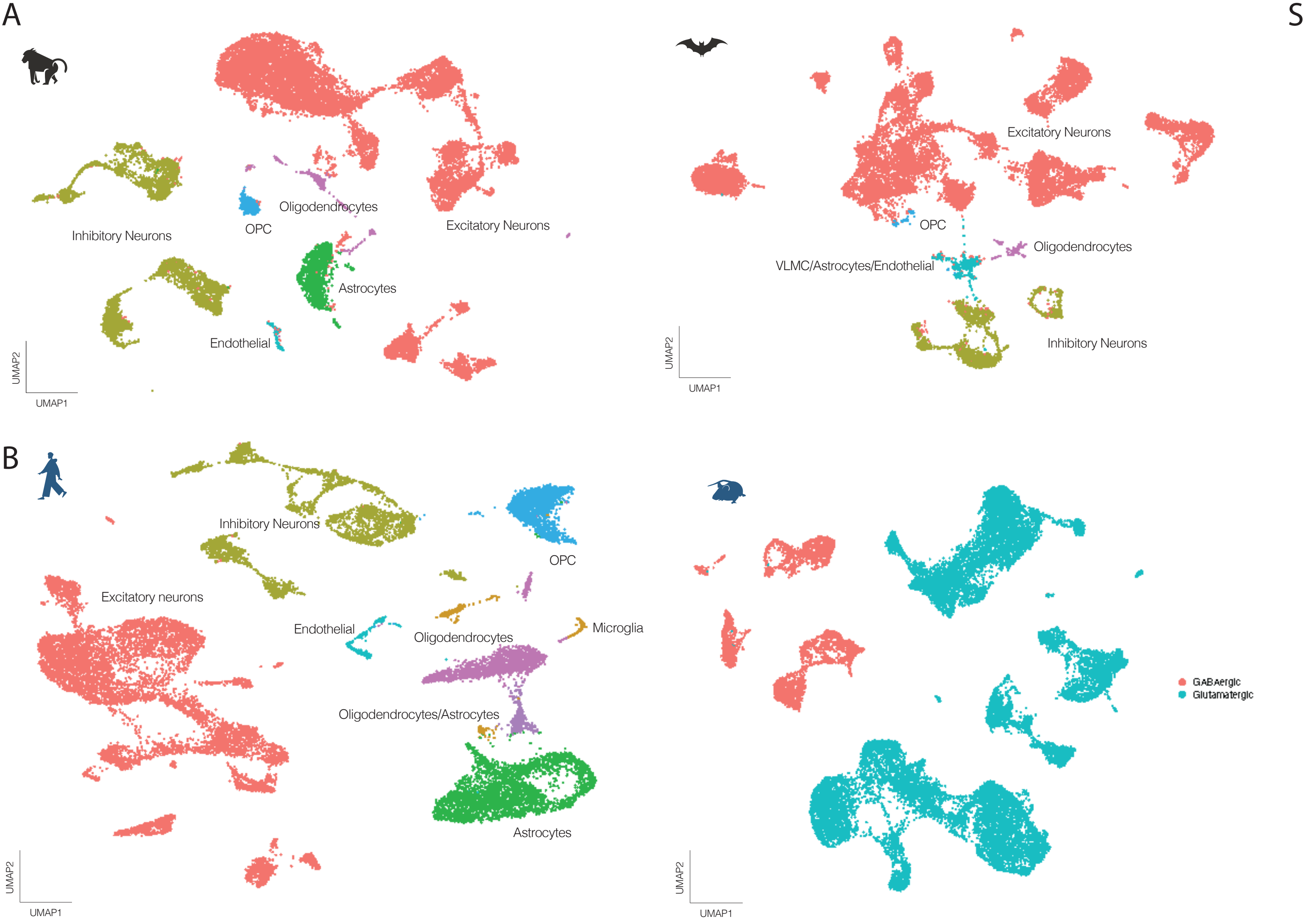
A
Inhibitory Neurons
OPC
Oligodendrocytes
Excitatory Neurons
Astrocytes
Endothelial
UMAP2
UMAP1
Excitatory Neurons
OPC
Oligodendrocytes
VLMC/Astrocytes/Endothelial
Inhibitory Neurons
UMAP2
UMAP1
B
Inhibitory Neurons
OPC
Excitatory neurons
Endothelial
Oligodendrocytes
Microglia
Oligodendrocytes/Astrocytes
Astrocytes
UMAP2
UMAP1
GABAergic
Glutamatergic
UMAP2
UMAP1

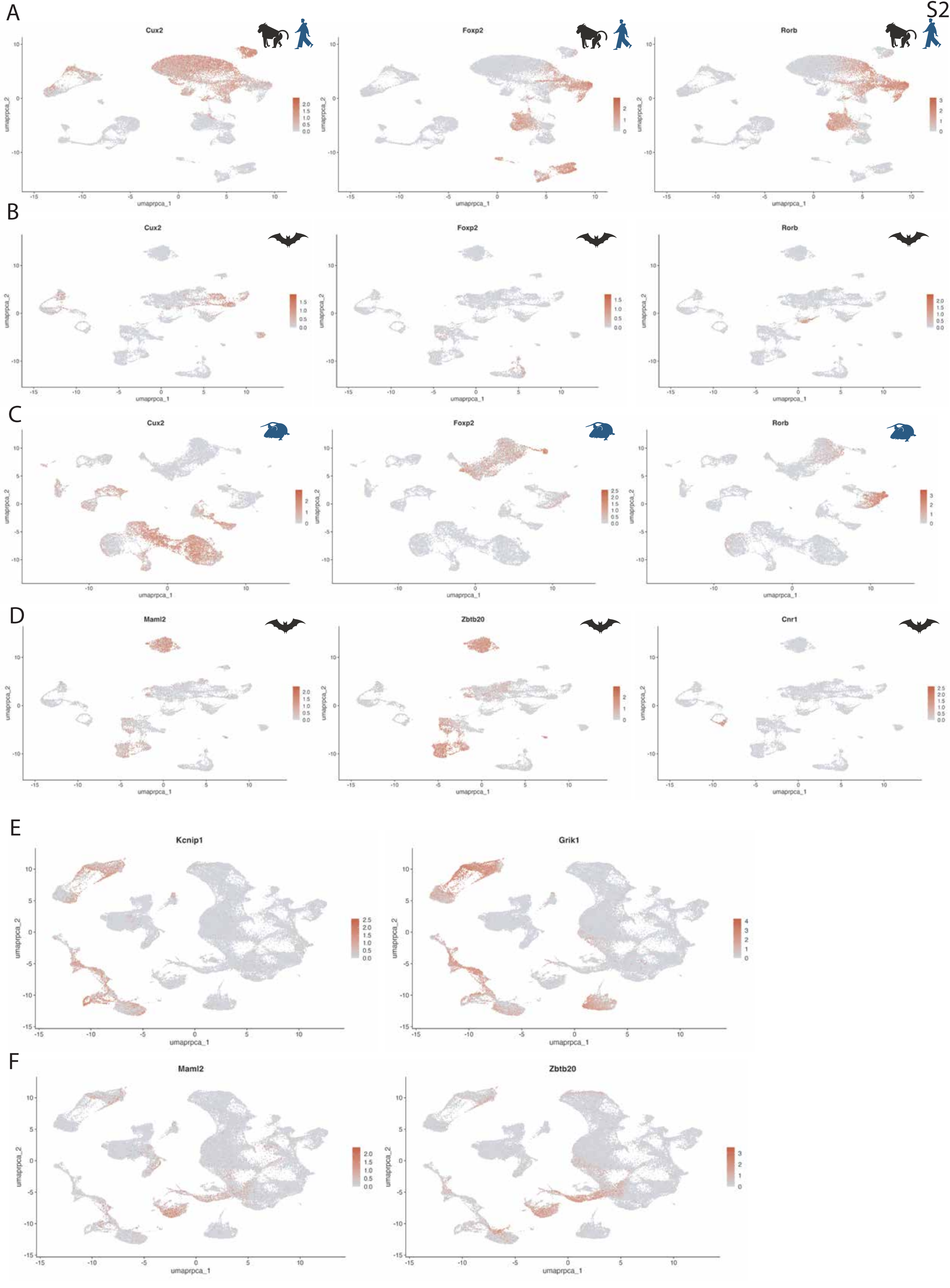
S2
A
Cux2
Foxp2
Rorb
B
Cux2
Foxp2
Rorb
C
Cux2
Foxp2
Rorb
D
Maml2
Zbtb20
Cnr1
E
Kcnip1
Grik1
F
Maml2
Zbtb20
umaprpca_1
umaprpca_2

S3

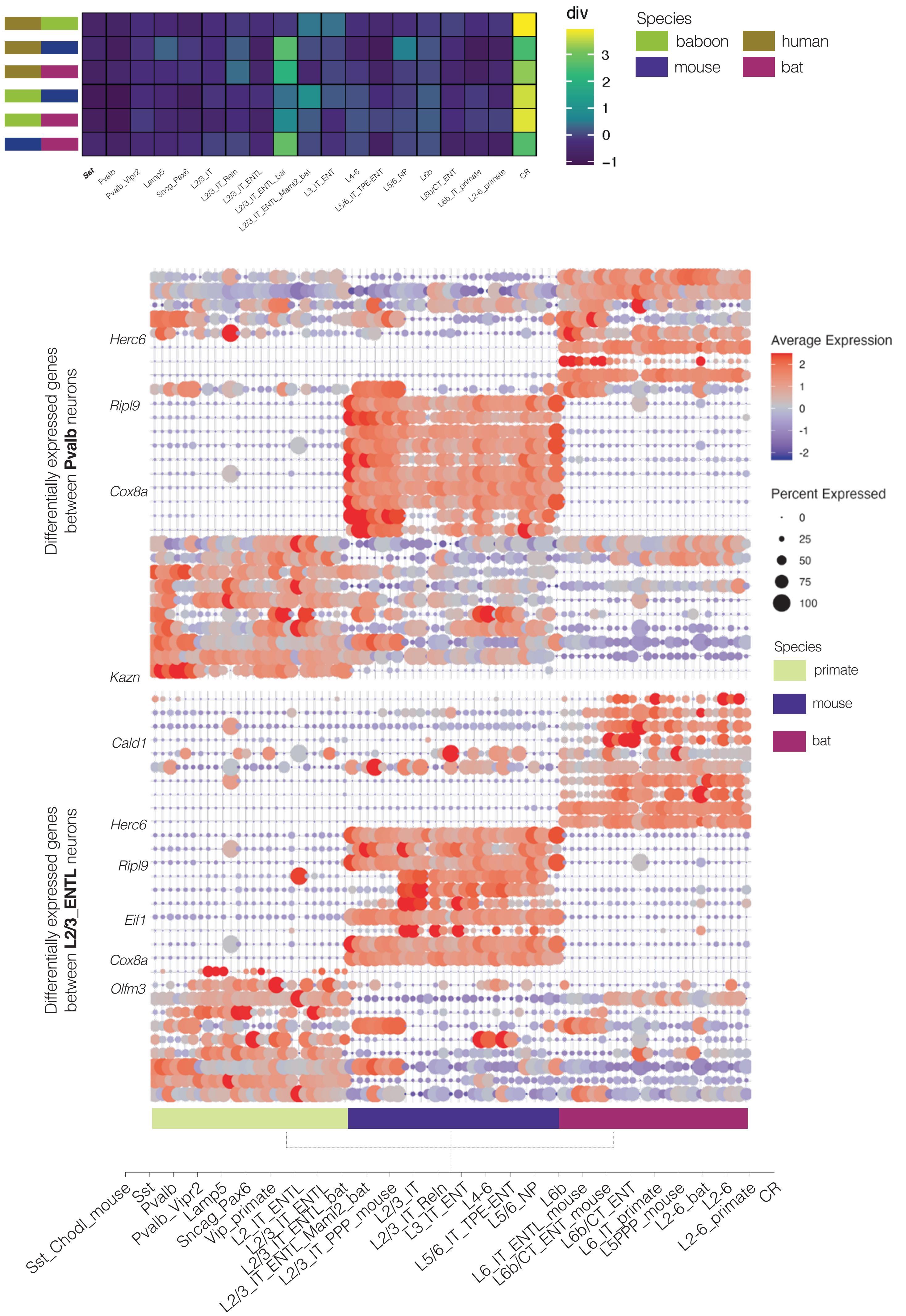

div
3
2
1
0
−1
Species
baboon
human
mouse
bat
Sst
Pvalb
Pvalb_Vipr2
Lamp5
Sncg_Pax6
L2/3_IT
L2/3_IT_Reln
L2/3_IT_ENTL
L2/3_IT_ENTL_bat
L2/3_IT_ENTL_Maml2_bat
L3_IT_ENT
L4-6
L5/6_IT_TPE-ENT
L5/6_NP
L6b
L6b/CT_ENT
L6b_IT_primate
L2-6_primate
CR
Differentially expressed genes between Pvalb neurons
Herc6
Ripl9
Cox8a
Kazn
Differentially expressed genes between L2/3_ENTL neurons
Cald1
Herc6
Ripl9
Eif1
Cox8a
Olfm3
Average Expression
2
1
0
-1
-2
Percent Expressed
0
25
50
75
100
Species
primate
mouse
bat
Sst_Chodl_mouse
Sst
Pvalb
Pvalb_Vipr2
Lamp5
Sncag_Pax6
Vip_primate
L2_IT_ENTL
L2/3_IT_ENTL
L2/3_IT_ENTL_bat
L2/3_IT_ENTL_Maml2_bat
L2/3_IT_PPP_mouse
L2/3_IT
L2/3_IT_Reln
L3_IT_ENT
L4-6
L5/6_IT_TPE-ENT
L5/6_NP
L6b
L6_IT_ENTL_mouse
L6b/CT_ENT_mouse
L6b/CT_ENT
L6_IT_primate
L5PPP_mouse
L2-6_bat
L2-6
L2-6_primate
CR

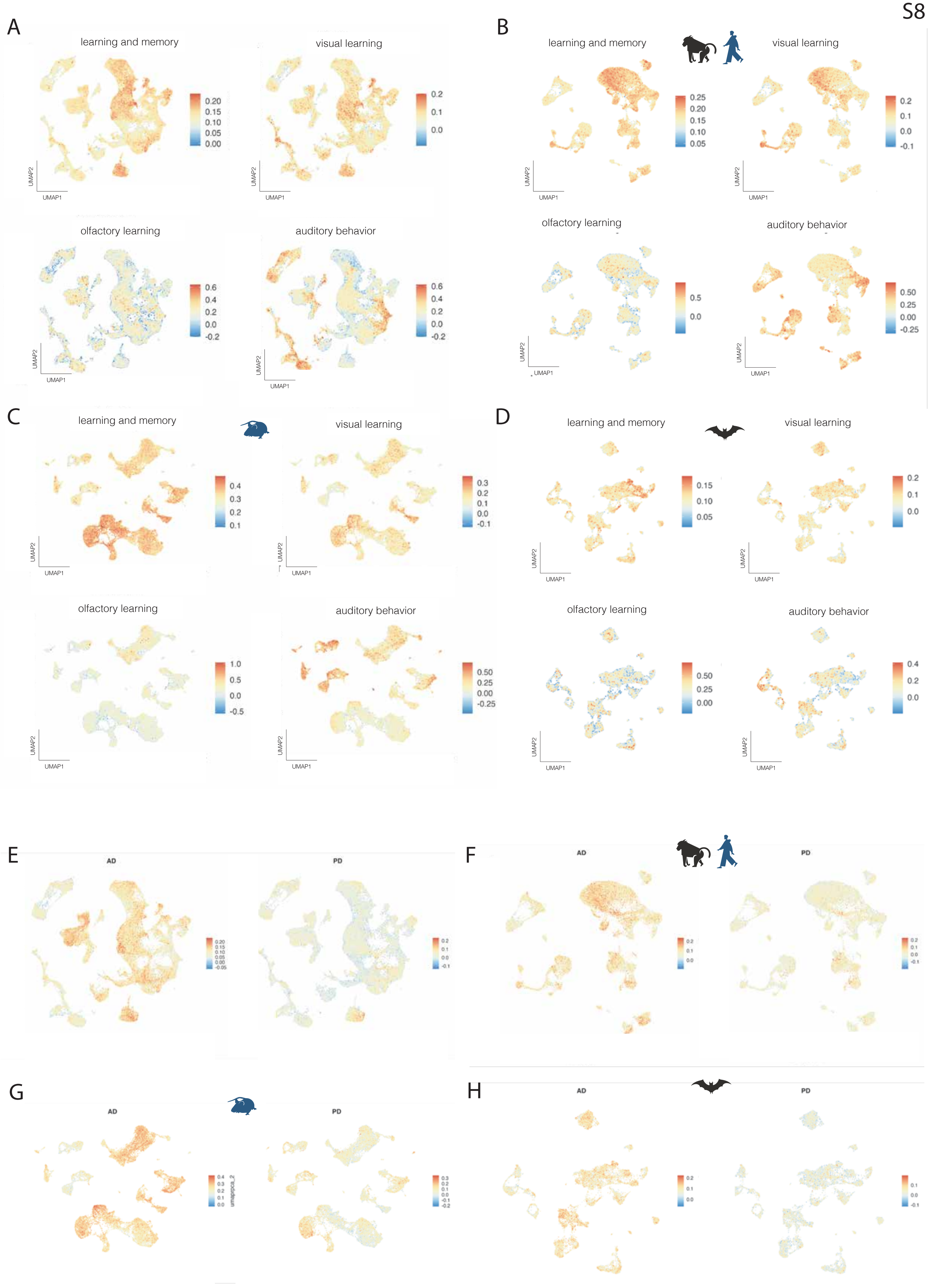
A
learning and memory
visual learning
olfactory learning
auditory behavior
B
learning and memory
visual learning
olfactory learning
auditory behavior
C
learning and memory
visual learning
olfactory learning
auditory behavior
D
learning and memory
visual learning
olfactory learning
auditory behavior
UMAP1
UMAP2
E
AD
PD
F
AD
PD
G
AD
PD
H
AD
PD

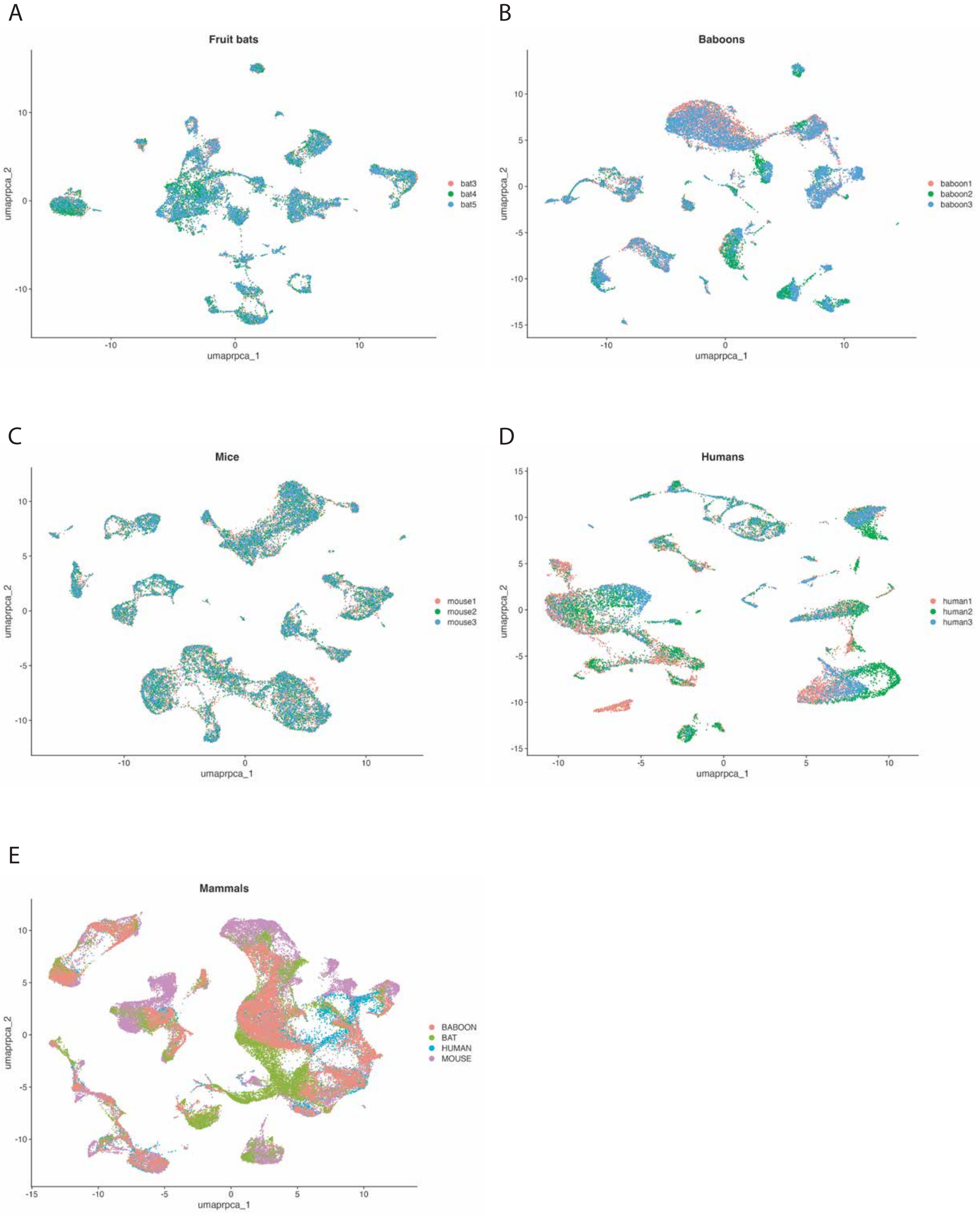
A
Fruit bats
bat3
bat4
bat5
umaprpca_1
umaprpca_2
B
Baboons
baboon1
baboon2
baboon3
umaprpca_1
umaprpca_2
C
Mice
mouse1
mouse2
mouse3
umaprpca_1
umaprpca_2
D
Humans
human1
human2
human3
umaprpca_1
umaprpca_2
E
Mammals
BABOON
BAT
HUMAN
MOUSE
umaprpca_1
umaprpca_2

A

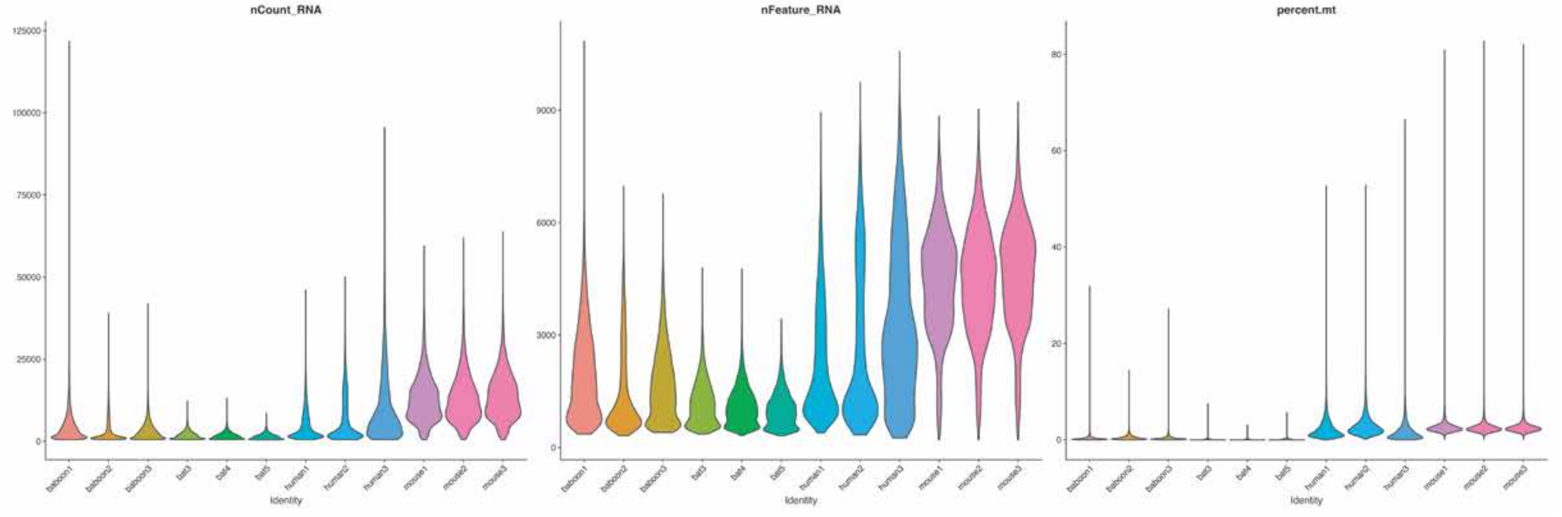
nCount_RNA
nFeature_RNA
percent.mt
baboon1
baboon2
baboon3
bat3
bat4
bat5
human1
human2
human3
mouse1
mouse2
mouse3
Identity

B

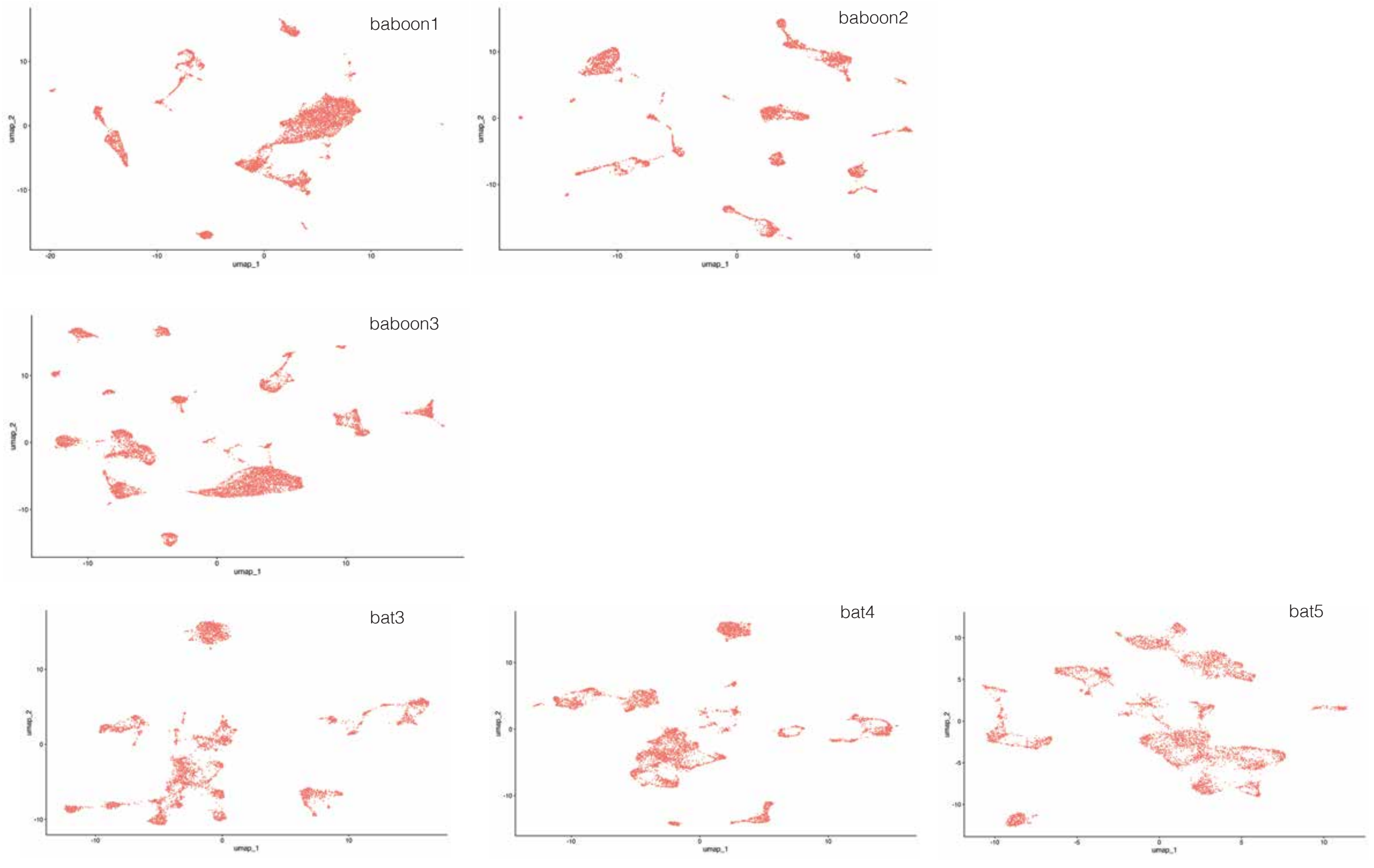
baboon1
baboon2
baboon3
bat3
bat4
bat5
umap_1
umap_2

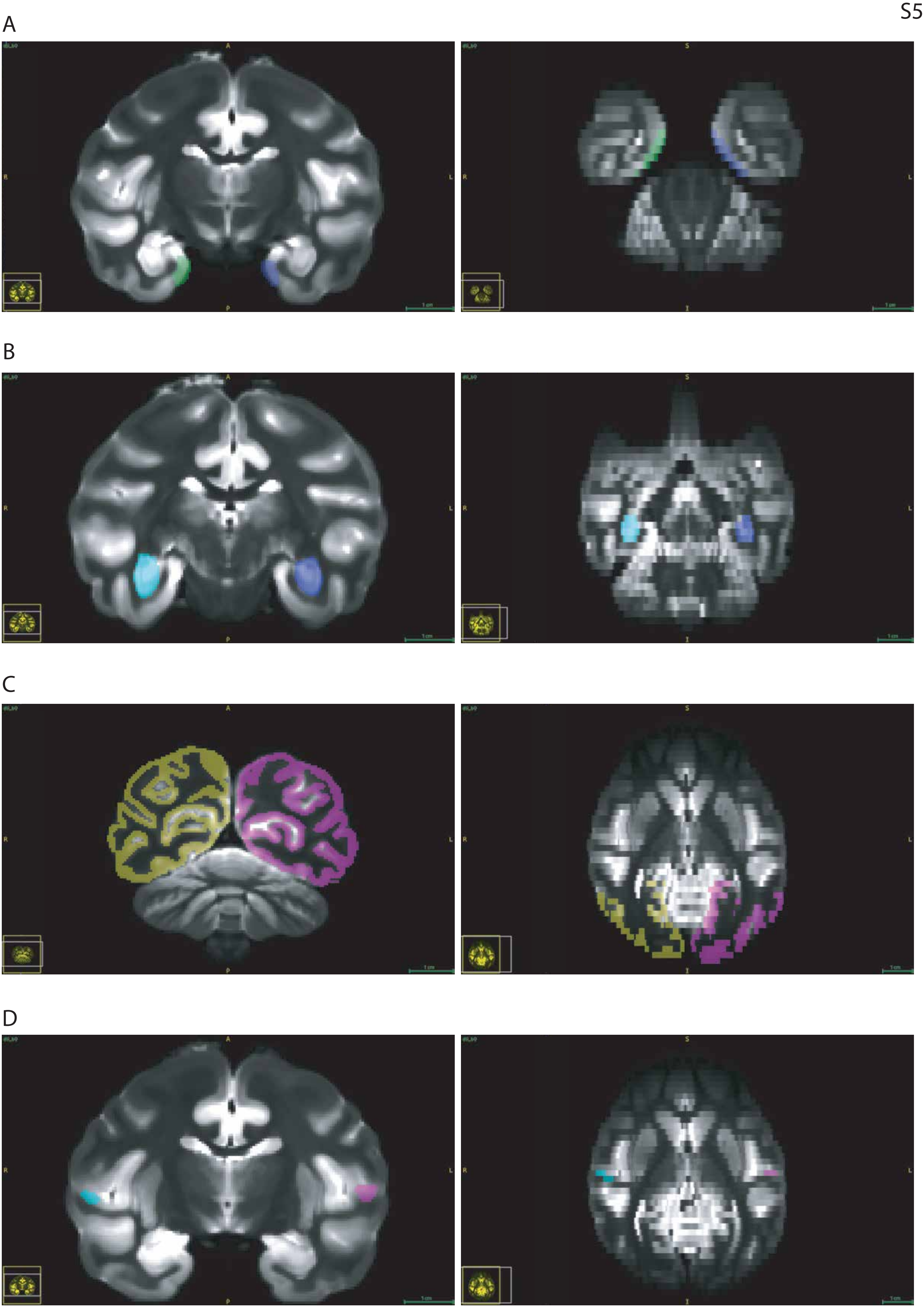
A
B
C
D

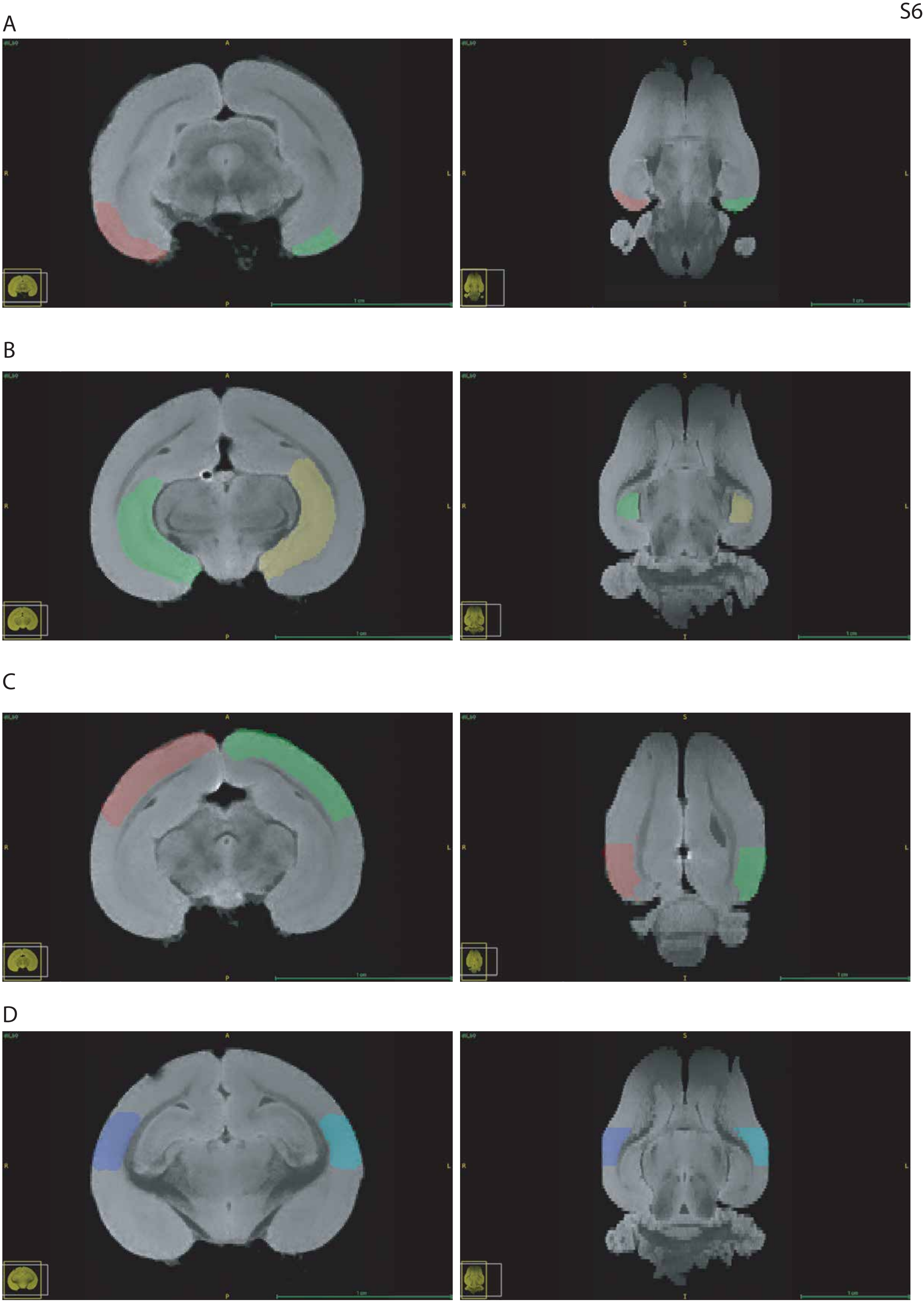
A
B
C
D

A

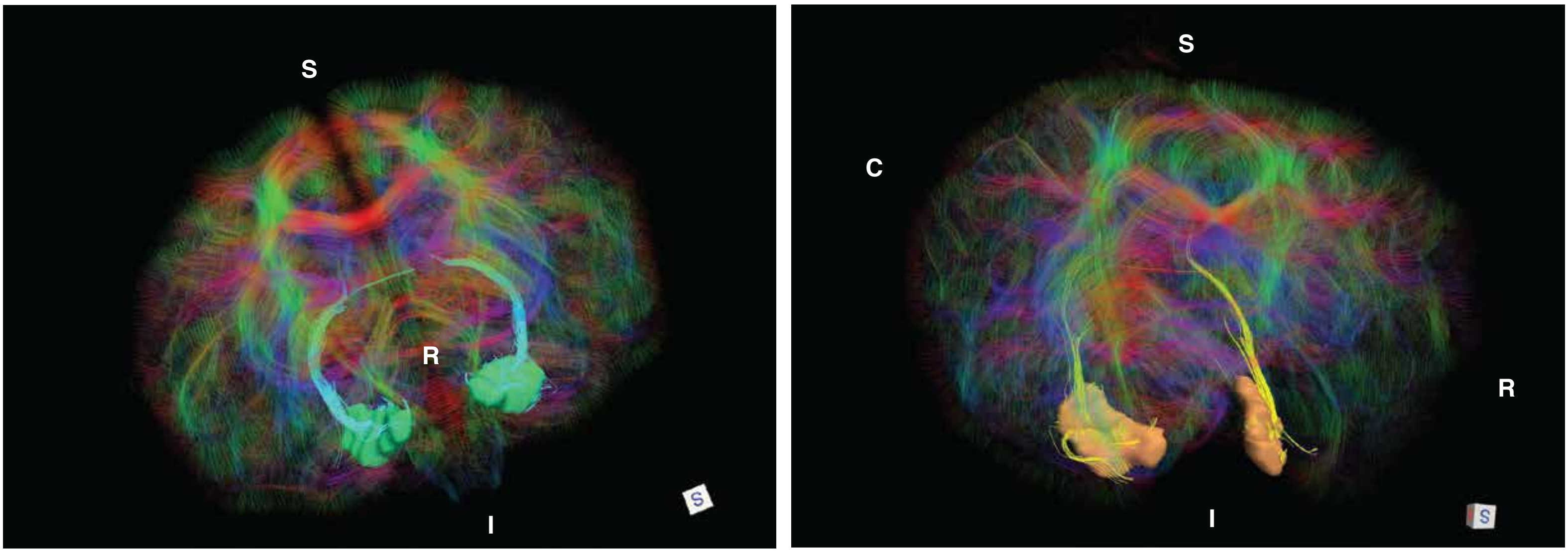


B

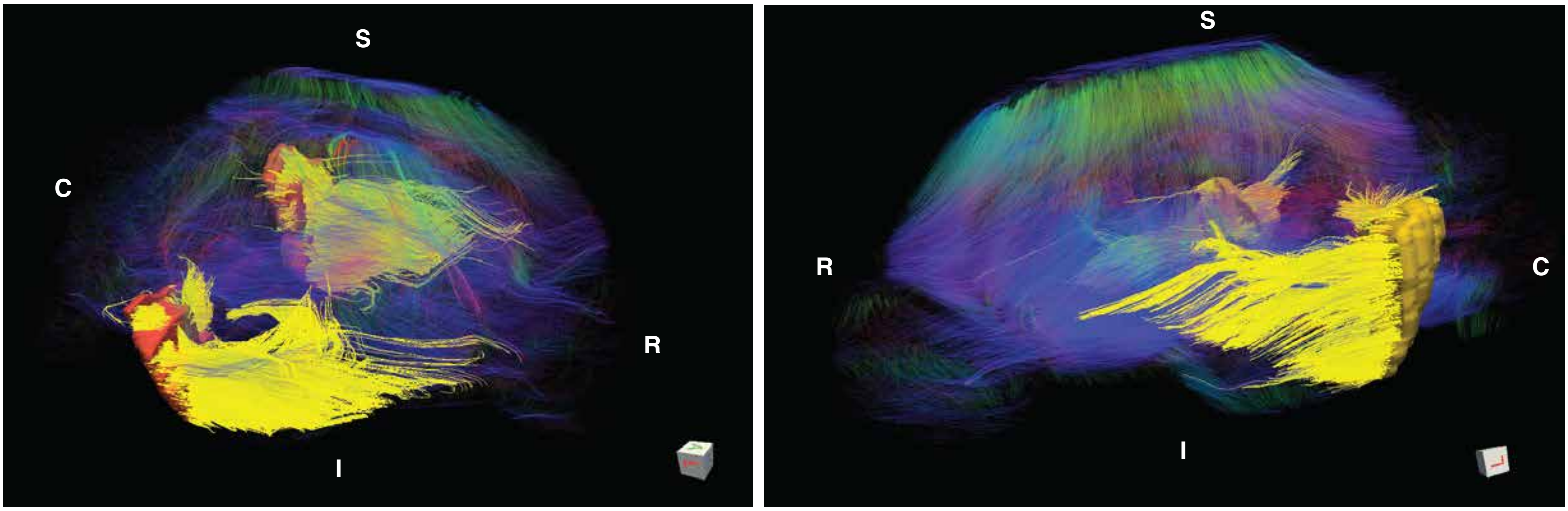

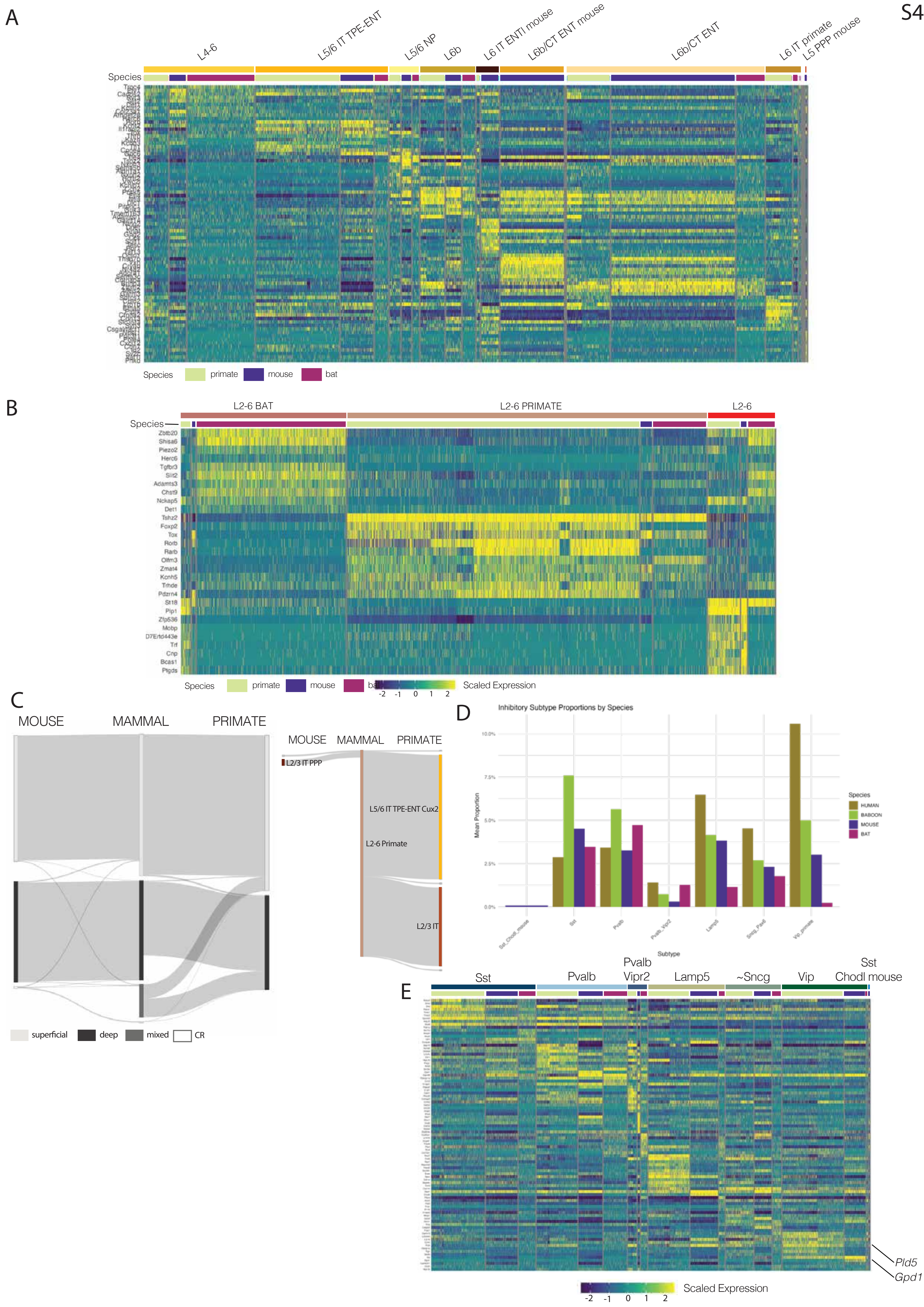
S4
A
L4-6
L5/6 IT TPE-ENT
L5/6 NP
L6b
L6 IT ENTl mouse
L6b/CT ENT mouse
L6b/CT ENT
L6 IT primate
L5 PPP mouse
Species
primate
mouse
bat
B
L2-6 BAT
L2-6 PRIMATE
L2-6
Zbtb20
Shisa6
Piezo2
Herc6
Tgfbr3
Slit2
Adamts3
Chst9
Nckap5
Det1
Tshz2
Foxp2
Tox
Rorb
Rarb
Olfm3
Zmat4
Kcnh5
Trhde
Pdzrn4
St18
Plp1
Zfp536
Mobp
D7Ertd443e
Trf
Cnp
Bcas1
Ptgds
Scaled Expression
C
MOUSE
MAMMAL
PRIMATE
L2/3 IT PPP
L5/6 IT TPE-ENT Cux2
L2-6 Primate
L2/3 IT
superficial
deep
mixed
CR
D
Inhibitory Subtype Proportions by Species
Mean Proportion
Subtype
Species
HUMAN
BABOON
MOUSE
BAT
E
Sst
Pvalb
Pvalb Vipr2
Lamp5
~Sncg
Vip
Sst Chodl mouse
Pld5
Gpd1
Scaled Expression

A

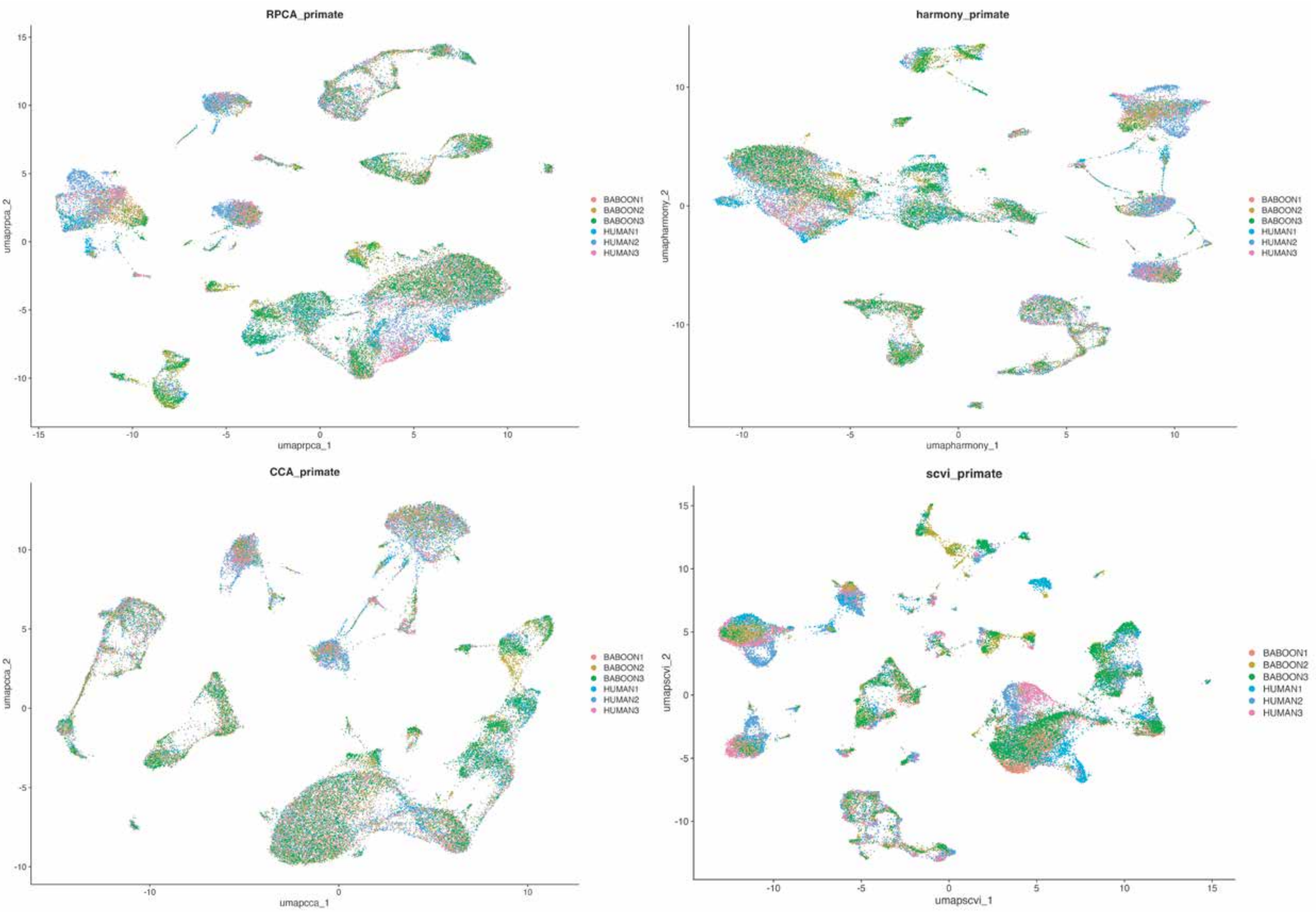
RPCA_primate
umaprpca_1
umaprpca_2
harmony_primate
umapharmony_1
umapharmony_2
CCA_primate
umapcca_1
umapcca_2
scvi_primate
umapscvi_1
umapscvi_2
BABOON1
BABOON2
BABOON3
HUMAN1
HUMAN2
HUMAN3

B

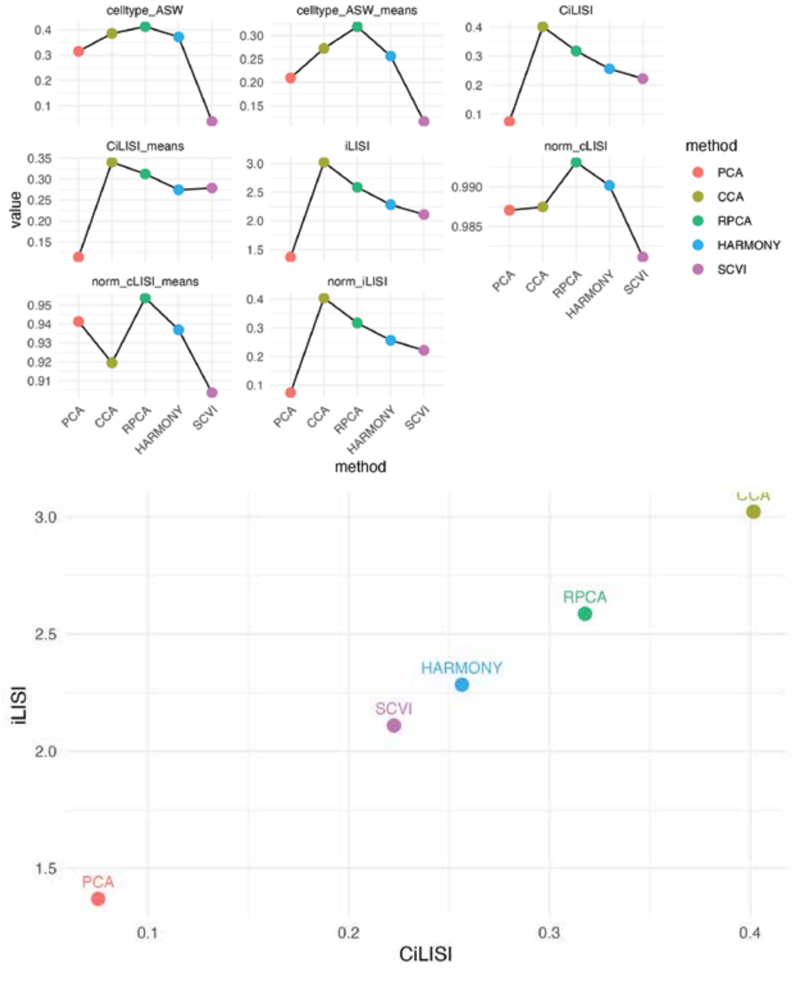
celltype_ASW
celltype_ASW_means
CiLISI
CiLISI_means
iLISI
norm_cLISI
norm_cLISI_means
norm_iLISI
value
method
PCA
CCA
RPCA
HARMONY
SCVI
iLISI
CiLISI

S9

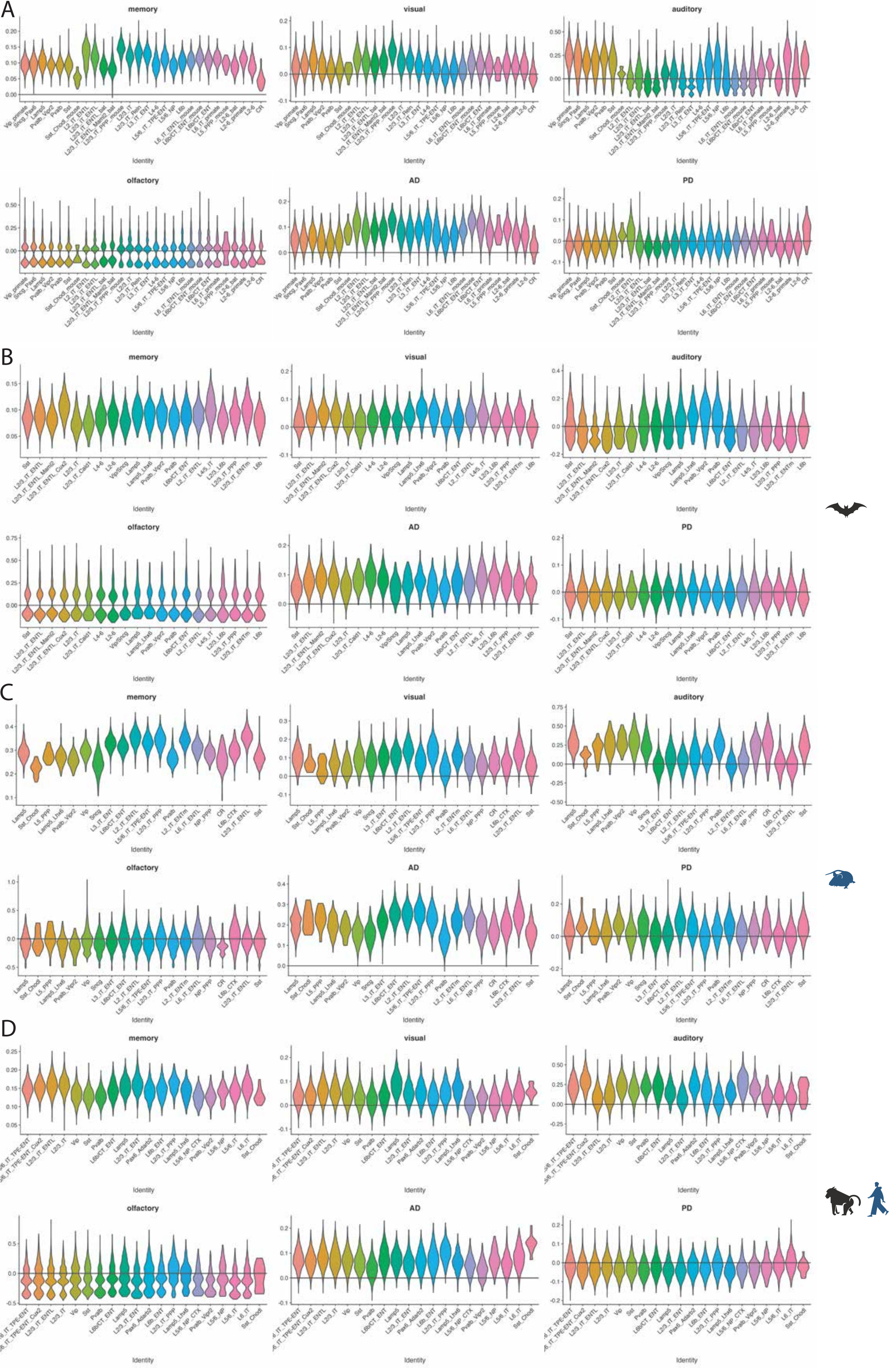
A
memory
visual
auditory
olfactory
AD
PD
Identity
B
memory
visual
auditory
olfactory
AD
PD
Identity
C
memory
visual
auditory
olfactory
AD
PD
Identity
D
memory
visual
auditory
olfactory
AD
PD
Identity

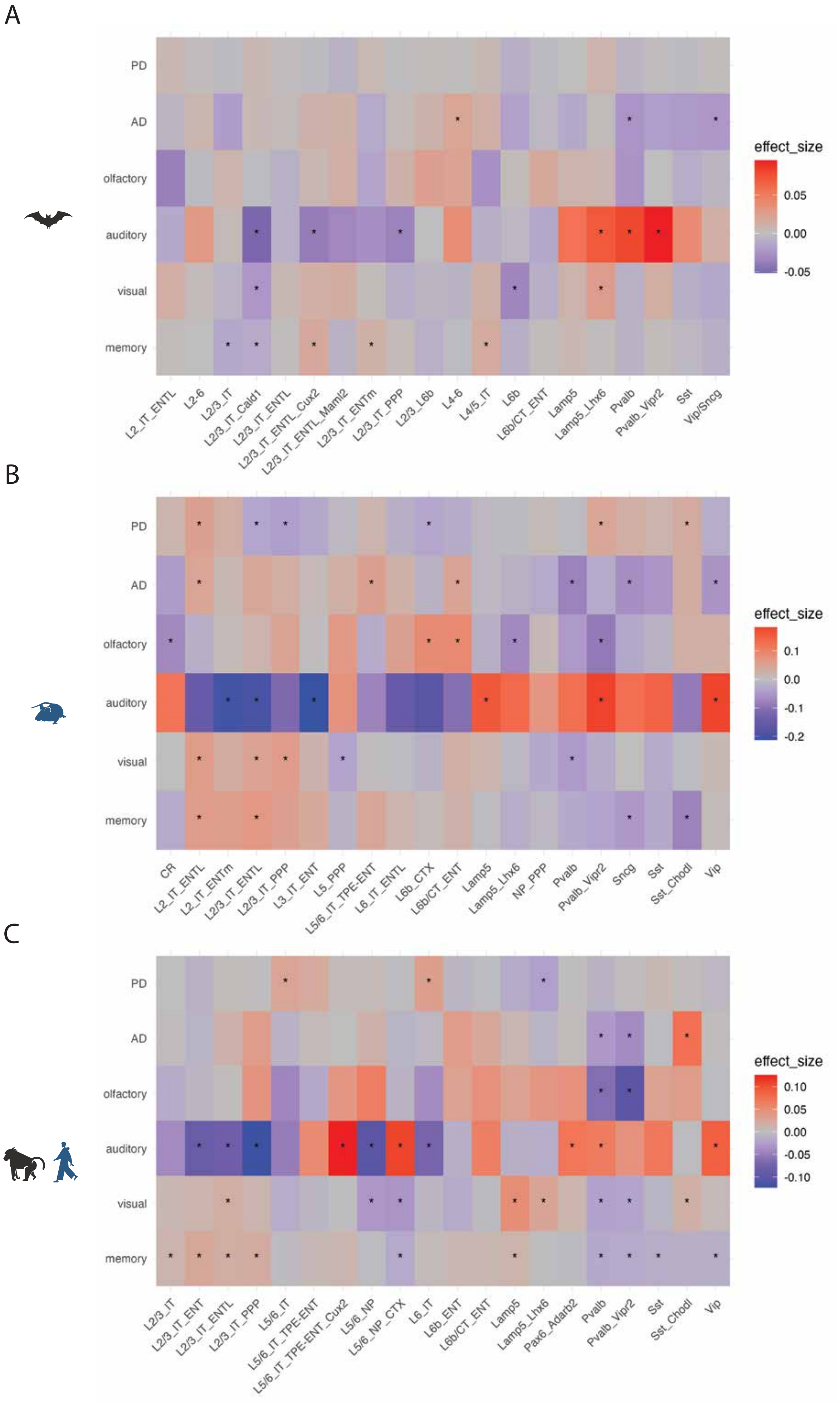
A
PD
AD
olfactory
auditory
visual
memory
L2_IT_ENTL
L2-6
L2/3_IT
L2/3_IT_Cald1
L2/3_IT_ENTL
L2/3_IT_ENTL_Cux2
L2/3_IT_ENTL_Maml2
L2/3_IT_ENTm
L2/3_IT_PPP
L2/3_L6b
L4-6
L4/5_IT
L6b
L6b/CT_ENT
Lamp5
Lamp5_Lhx6
Pvalb
Pvalb_Vipr2
Sst
Vip/Sncg
effect_size
0.05
0.00
-0.05
B
PD
AD
olfactory
auditory
visual
memory
CR
L2_IT_ENTL
L2_IT_ENTm
L2/3_IT_ENTL
L2/3_IT_PPP
L3_IT_ENT
L5_PPP
L5/6_IT_TPE-ENT
L6_IT_ENTL
L6b_CTX
L6b/CT_ENT
Lamp5
Lamp5_Lhx6
NP_PPP
Pvalb
Pvalb_Vipr2
Sncg
Sst
Sst_Chodl
Vip
effect_size
0.1
0.0
-0.1
-0.2
C
PD
AD
olfactory
auditory
visual
memory
L2/3_IT
L2/3_IT_ENT
L2/3_IT_ENTL
L2/3_IT_PPP
L5/6_IT
L5/6_IT_TPE-ENT
L5/6_IT_TPE-ENT_Cux2
L5/6_NP
L5/6_NP_CTX
L6_IT
L6b_ENT
L6b/CT_ENT
Lamp5
Lamp5_Lhx6
Pax6_Adarb2
Pvalb
Pvalb_Vipr2
Sst
Sst_Chodl
Vip
effect_size
0.10
0.05
0.00
-0.05
-0.10
